# Improved genetically encoded fluorescent biosensors for monitoring of intra- and extracellular L-lactate

**DOI:** 10.1101/2022.12.27.522013

**Authors:** Yusuke Nasu, Abhi Aggarwal, Giang N. T. Le, Yuki Kamijo, Marc Boisvert, Marie-Eve Paquet, Mikhail Drobizhev, Kaspar Podgorski, Robert E. Campbell

## Abstract

l-Lactate is increasingly appreciated as a key metabolite and signaling molecule in mammals. To enable investigations of both the inter- and intra-cellular dynamics of l-Lactate, we develop a second-generation green fluorescent extracellular l-Lactate biosensor, designated eLACCO2.1, and a red fluorescent intracellular l-Lactate biosensor, designated R-iLACCO1. Compared to the first-generation eLACCO1.1 (Δ*F*/*F* = 1.5 in cultured neurons), eLACCO2.1 exhibits better membrane localization and fluorescence response (Δ*F*/*F* = 8.1 in cultured neurons) with faster response kinetics to extracellular l-Lactate on the surface of live mammalian cells. R-iLACCO1 and its affinity variants exhibit large fluorescence responses to changes in l-Lactate concentration *in vitro* (Δ*F*/*F* = 15 to 22) and in live mammalian cells (Δ*F*/*F* = 5.5 to 11). We demonstrate that these biosensors enable cellular-resolution imaging of extracellular and intracellular l-Lactate in cultured mammalian cells.

## Introduction

l-Lactate was once considered a waste by-product of glucose metabolism^1^. However, growing evidence suggests that l-Lactate plays a variety of important roles as both an energy fuel and a signaling molecule in the nervous system^2^, tumor microenvironment^3^, gut microbiome^4^, and immune system^5^. These roles have impacts on physiological and pathological processes over scales and environments ranging from subcellular^6^, to intercellular^2, 7^, to interorgan^8^.

Investigations of the emerging roles of l-Lactate in cells and tissues would be facilitated by a set of genetically encoded fluorescent biosensors that could enable high resolution spatially and temporally resolved imaging of l-Lactate in the extracellular and intracellular environment. As steps towards this end, we previously developed two classes of genetically encoded green fluorescent biosensors: <u>e</u>LACCO1.1 for <u>e</u>xtracellular l-Lactate^9^ and iLACCO1 for intracellular l-Lactate^10^. In addition to our LACCO series, others have developed cyan or green fluorescent genetically encoded biosensors for intracellular l-Lactate^11–17^. However, the first-in-class eLACCO1.1 suffers from aggregation and limited fluorescence response in mammalian cells. Another limitation of the current set of l-Lactate biosensors is that no red fluorescent biosensors for intracellular l-Lactate have yet been reported, hindering multiplexed l-Lactate imaging with blue-light activated optogenetic actuators or other green fluorescent biosensors.

In this work, we have developed a spectrally and functionally orthogonal pair of high-performance genetically encoded biosensors: the second-generation green fluorescent eLACCO2.1 for extracellular l-Lactate and the first-generation red fluorescent R-iLACCO1 for intracellular l-Lactate (**Fig. 1a,b**). We confirm that eLACCO2.1 and R-iLACCO1 enable cellular resolution imaging of extracellular and intracellular l-Lactate, respectively, in cultured mammalian cells.

**Fig. 1.**
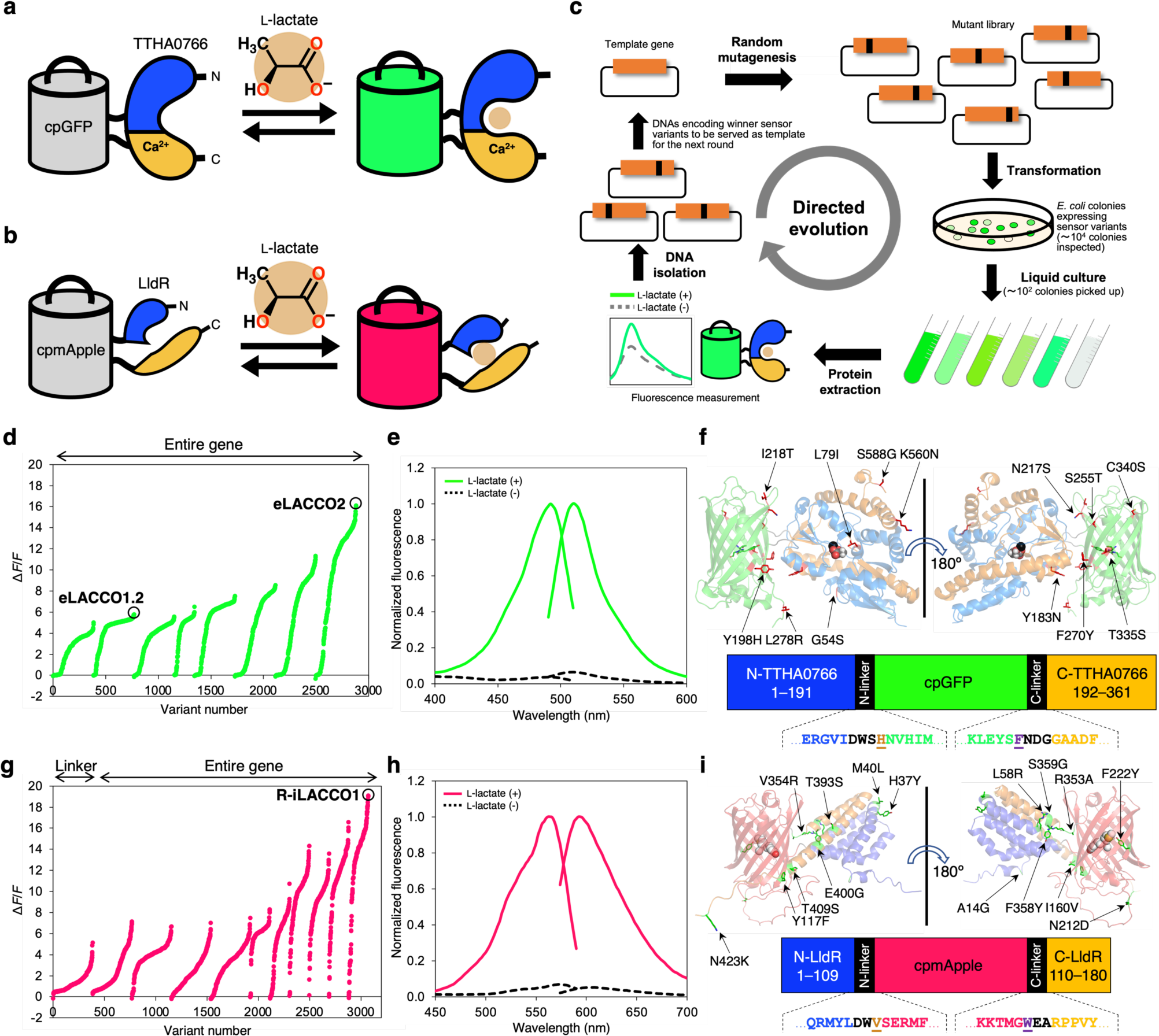
Development of eLACCO2.1 and R-iLACCO1. (**a**) Schematic representation of TTHA0766-based eLACCO and its mechanism of response to l-Lactate. (**b**) Schematic representation of LldR-based R-iLACCO and its mechanism of response to l-Lactate. (**c**) Schematic of directed evolution workflow. Specific sites (i.e., the linkers) or the entire gene of template l-Lactate biosensor were randomly mutated and the resulting mutant library was used to transform *E. coli*. Bright colonies were picked and cultured, and then proteins were extracted to examine Δ*F*/*F* upon addition of 10 mM l-Lactate. A mixture of the variants with the highest Δ*F*/*F* was used as the template for the next round. (**d**) Δ*F*/*F* rank plot representing all crude proteins tested during the directed evolution of eLACCO. For each round, tested variants are ranked from lowest to highest Δ*F*/*F* value from left to right. (**e**) Excitation and emission spectra of purified eLACCO2.1 in the presence (10 mM) and absence of l-Lactate. (**f**) Overall representation of the eLACCO1 crystal structure (PDB 7E9Y) with the position of mutations indicated. l-Lactate and Ca^2+^ (black) are shown in a sphere representation. In the primary structure of eLACCO2.1 (bottom), linker regions are shown in black and the two “gate post” residues^33^ in cpGFP are highlighted in dark orange (His195) and purple (Phe437). (**g**) Δ*F*/*F* rank plot representing all crude proteins tested during the directed evolution of R-iLACCO. (**h**) Excitation and emission spectra of purified R-iLACCO1 in the presence (10 mM) and absence of l-Lactate. (**i**) Overall representation of the R-iLACCO1 model structure^37^ with the position of mutations indicated. Unmatured chromophore is shown in a sphere representation. In the primary structure of R-iLACCO1 (bottom), linker regions are shown in black and the two “gate post” residues^33^ in cpmApple are highlighted in dark orange (Val112) and purple (Trp351).

## Results

### Development of a second-generation green fluorescent l-Lactate biosensor, eLACCO2.1

Before embarking on an effort to develop an improved second-generation green fluorescent extracellular l-Lactate biosensor, we first considered which of the previously reported eLACCO variants would be the most promising starting point. We identified eLACCO0.9 as a promising candidate due to the fact that it exhibits a faster fluorescence response to l-Lactate than eLACCO1 (**Supplementary Fig. 1**). To develop variants of eLACCO0.9 with larger change in fluorescence intensity (Δ*F*/*F* = (*F*_max_ − *F*_min_)/ *F*_min_) upon l-Lactate treatment, we performed directed evolution with screening of crude protein extracts for Δ*F*/*F* (**Fig. 1c**). The first two rounds of evolution used libraries created by random mutagenesis of the entire gene of eLACCO0.9 (Δ*F*/*F* = 3.9) and resulted in the identification of the eLACCO1.2 variant with Δ*F*/*F* of 5.8 (**Fig. 1d**). eLACCO1.2 contains two cysteine residues (Cys340 and Cys362), both of which are located in cpGFP. To avoid the potential formation of disulfide bond between eLACCO proteins^18^, we introduced the Cys340Ser mutation into eLACCO1.2 to produce eLACCO1.3 (Δ*F*/*F* = 4.8, crude protein extract). Incorporation of Cys362Val into eLACCO1.3 abrogated the lactate-dependent response (Δ*F*/*F* = 0.8, crude protein extract) and therefore this variant (eLACCO1.3 Cys362Val) was not further pursued. To further improve the Δ*F*/*F*, we continued to perform directed evolution starting from eLACCO1.3. Six additional rounds of directed evolution led to the eLACCO2 variant with Δ*F*/*F* of 16 (**Fig. 1d**).

Characterization of purified eLACCO2 revealed that it has an affinity for l-Lactate (apparent *K*_d_ ∼ 280 μM) that is much higher than typical extracellular concentrations, which are in the millimolar range (**Supplementary Fig. 2**). To tune the affinity of eLACCO2 for imaging of extracellular l-Lactate, we decreased its affinity using mutagenesis of residues that interact with l-Lactate in the TTHA0766 lactate-binding pocket. The crystal structure of eLACCO1 (PDB 7E9Y) had revealed that the side chains of Tyr80, Phe77, and Leu79 line the l-Lactate-binding site^9^. We tested crude protein extracts of a series of ten conservative mutants (Phe77Ala, Phe77Ile, Phe77Leu, Phe77Val, Phe77Tyr, Leu79Ala, Leu79Phe, Leu79Ile, Leu79Val, and Tyr80Phe) that we expected to potentially have a modest effect on l-Lactate binding affinity (**Supplementary Fig. 2b**). Of the ten variants investigated, eLACCO2 Leu79Ile exhibited a lower affinity (apparent *K*_d_ = 1.9 mM) with a minimal decrease in Δ*F*/*F* compared to eLACCO2 (**Supplementary Fig. 2b,c**). The Tyr80Phe mutation substantially decreased the fluorescence response to l-Lactate and was not further pursued (**Supplementary Fig. 2c**). eLACCO2 Leu79Ile, designated eLACCO2.1, has a Δ*F*/*F* of 14 in purified protein, which is 3.5-fold higher than the first-generation eLACCO1.1 (**Fig. 1e** and **Supplementary Table 1**), and has 13 mutations relative to eLACCO0.9 (**Fig. 1f** and **Supplementary Fig. 3**). A non-responsive control biosensor, designated deLACCO1, was engineered by incorporating the Asp444Asn mutation into eLACCO2 to abolish l-Lactate binding and subsequently performing three rounds of directed evolution to improve fluorescence brightness (**Supplementary Figs. 3–5**).

### Development of a red fluorescent biosensor for intracellular l-Lactate, R-iLACCO1

To construct an initial prototype red fluorescent l-Lactate biosensor based on LldR transcriptional regulator protein (**Fig. 1b**), we inserted the circularly permuted red fluorescent protein (cpmApple), derived from the TTHA0766-based red fluorescent l-Lactate biosensor R-eLACCO1 (ref. 19), into the l-Lactate binding domain (LBD) of the *Escherichia coli* LldR at position 109 (**Supplementary Fig. 6a**)^10^. Relative to Ca^2+^-requiring TTHA0766, LldR is a preferred binding domain for an intracellular l-Lactate biosensor because LldR has no Ca^2+^ binding site and has been demonstrated to be functional in the cytosol where Ca^2+^ concentration is low (∼ 0.1–1 μM)^10, 11^. The resulting variant, designated R-iLACCO0.1, exhibited dim fluorescence and only a slight increase in fluorescence intensity (Δ*F*/*F* = 0.7, crude protein extract) upon treatment with l-Lactate (**Supplementary Fig. 6b**). To improve the performance of R-iLACCO0.1, we optimized the lengths of linkers that connect cpmApple and LldR, and found that a variant (designated R-iLACCO0.2) with two amino acids of the C-terminal linker deleted had substantially improved brightness, while retaining a comparable fluorescence response to l-Lactate (Δ*F*/*F* = 0.6, crude protein extract) (**Supplementary Fig. 6c,d**). To develop variants of R-iLACCO0.2 with larger Δ*F*/*F*, we performed one round of the length-optimized C-terminal linker optimization, followed by ten rounds of directed evolution by random mutagenesis of the whole gene, with screening for l-Lactate-dependent change in fluorescence intensity (**Fig. 1c,g**). This effort ultimately produced the R-iLACCO1 with Δ*F*/*F* of 20 (purified protein, **Fig. 1h**). R-iLACCO1 contains a total of 16 mutations relative to R-iLACCO0.1 (**Fig. 1i** and **Supplementary Fig. 7**).

The wide range of physiological concentrations of intracellular l-Lactate inspired us to engineer lower affinity variants of R-iLACCO1. To this end, in the final round of directed evolution we screened all randomly picked variants in the absence of l-Lactate, at 0.5 mM l-Lactate, and at 10 mM l-Lactate, as a way of assessing their apparent *K*_d_ values. This effort led us to identify two lower affinity variants, designated R-iLACCO1.1 and R-iLACCO1.2 (**Supplementary Fig. 8**). To engineer a non-responsive control biosensor, designated R-diLACCO1, we incorporated the Asp69Asn mutation into R-iLACCO1 and then performed two rounds of directed evolution to improve fluorescence brightness (**Supplementary Figs. 4b, 5c**, and **7**).

### *In vitro* characterization of eLACCO2.1

We characterized the photophysical and biochemical properties of purified eLACCO2.1 in a soluble form (**Fig. 2a–f**, **Table 1,** and **Supplementary Table 1**). In the presence of l-Lactate, eLACCO2.1 exhibits absorbance peaks at 398 nm and 494 nm (**Fig. 2a**), corresponding to the neutral (protonated) and the anionic (deprotonated) forms of chromophore, respectively. Addition of l-Lactate changes these two absorbance peaks in a ratiometric manner, consistent with a shift in the equilibrium from a neutral form-dominant state to an anionic form-dominant state (**Fig. 2a** and **Table 1**). eLACCO2.1 in the presence of l-Lactate has an excitation maximum at 495 nm and the emission maximum is 509 nm (**Fig. 1e**). The molecular brightness of eLACCO2.1 in the l-Lactate-bound state is 137% that of eLACCO1.1 and 140% and 100% that of the jGCaMP7s (+Ca^2+^) and EGFP, respectively (**Table 1**)^20^. The two-photon excitation maximum of l-Lactate bound eLACCO2.1 is 932 nm with a brightness of *F*_2_ = 20 GM that is 25% higher than l-Lactate-bound eLACCO1.1 (**Fig. 2c** and **Table 1**). The Δ*F*_2_/*F*_2_ value in the 932–1000 nm wavelength ranges from 20–25, which is a 2–3-fold higher than eLACCO1.1 (**Fig. 2c** and **Table 1**).

**Fig. 2.**
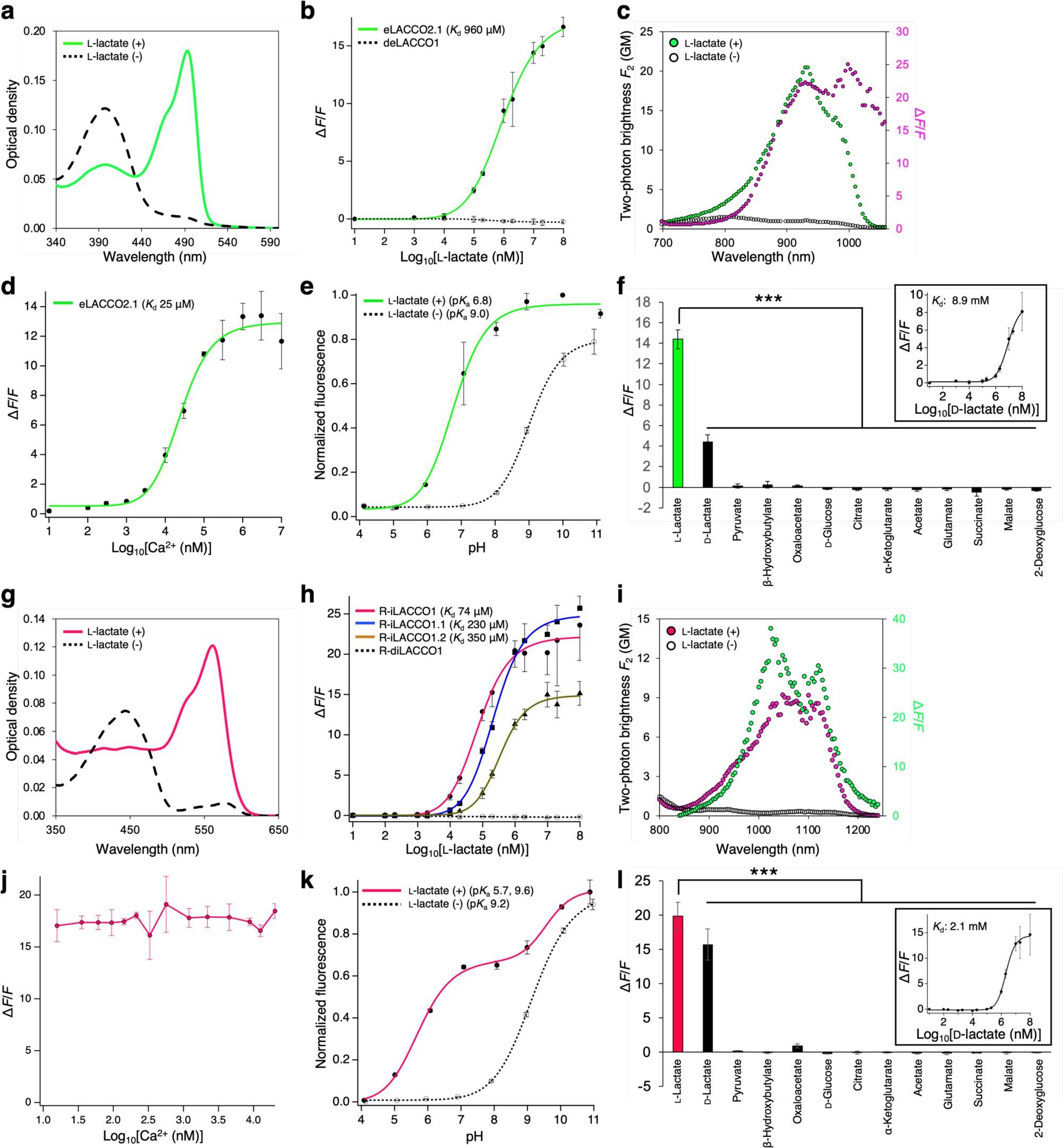
*In vitro* characterization of eLACCO2.1 and R-iLACCO1. (**a**) Absorbance spectra of eLACCO2.1 in the presence (10 mM) and absence of l-Lactate. (**b**) Dose-response curves of eLACCO2.1 and deLACCO1 for l-Lactate. *n* = 3 experimental triplicates (mean ± s.d.). (**c**) Two-photon excitation spectra of eLACCO2.1 in the presence (10 mM) and absence of l-Lactate. Δ*F*/*F* is represented in the magenta plots. GM, Goeppert-Mayer units. (**d**) Dose-response curve of eLACCO2.1 as a function of Ca^2+^ in the presence (100 mM) and absence of l-Lactate. *n* = 3 experimental triplicates (mean ± s.d.). (**e**) pH titration curves of eLACCO2.1 in the presence (100 mM) and absence of l-Lactate. *n* = 3 experimental triplicates (mean ± s.d.). (**f**) Pharmacological specificity of eLACCO2.1. Concentration of each metabolite is 10 mM. Inset is a dose-response curve of eLACCO2.1 for d-lactate. *n* = 3 experimental triplicates (mean ± s.d.). Statistical analysis was performed using one-way analysis of variance (ANOVA) with the Dunnett’s post hoc tests. \*\*\**p* < 0.0001. (**g**) Absorbance spectra of R-iLACCO1 in the presence (10 mM) and absence of l-Lactate. (**h**) Dose-response curves of R-iLACCO1 and its affinity variants for l-Lactate. *n* = 3 experimental triplicates (mean ± s.d.). (**i**) Two-photon excitation spectra of R-iLACCO1 in the presence (10 mM) and absence of l-Lactate. Δ*F*/*F* is represented in the green plots. GM, Goeppert-Mayer units. (**j**) Dose-response curve of R-iLACCO1 as a function of Ca^2+^ in the presence (100 mM) and absence of l-Lactate. *n* = 3 experimental triplicates (mean ± s.d.). (**k**) pH titration curves of R-iLACCO1 in the presence (100 mM) and absence of l-Lactate. *n* = 3 experimental triplicates (mean ± s.d.). (**l**) Pharmacological specificity of R-iLACCO1. Concentration of each metabolite is 10 mM. Inset is a dose-response curve of R-iLACCO1 for d-lactate. *n* = 3 experimental triplicates (mean ± s.d.). Statistical analysis was performed using one-way ANOVA with the Dunnett’s post hoc tests. \*\*\**p* < 0.0001.

**Table 1.**
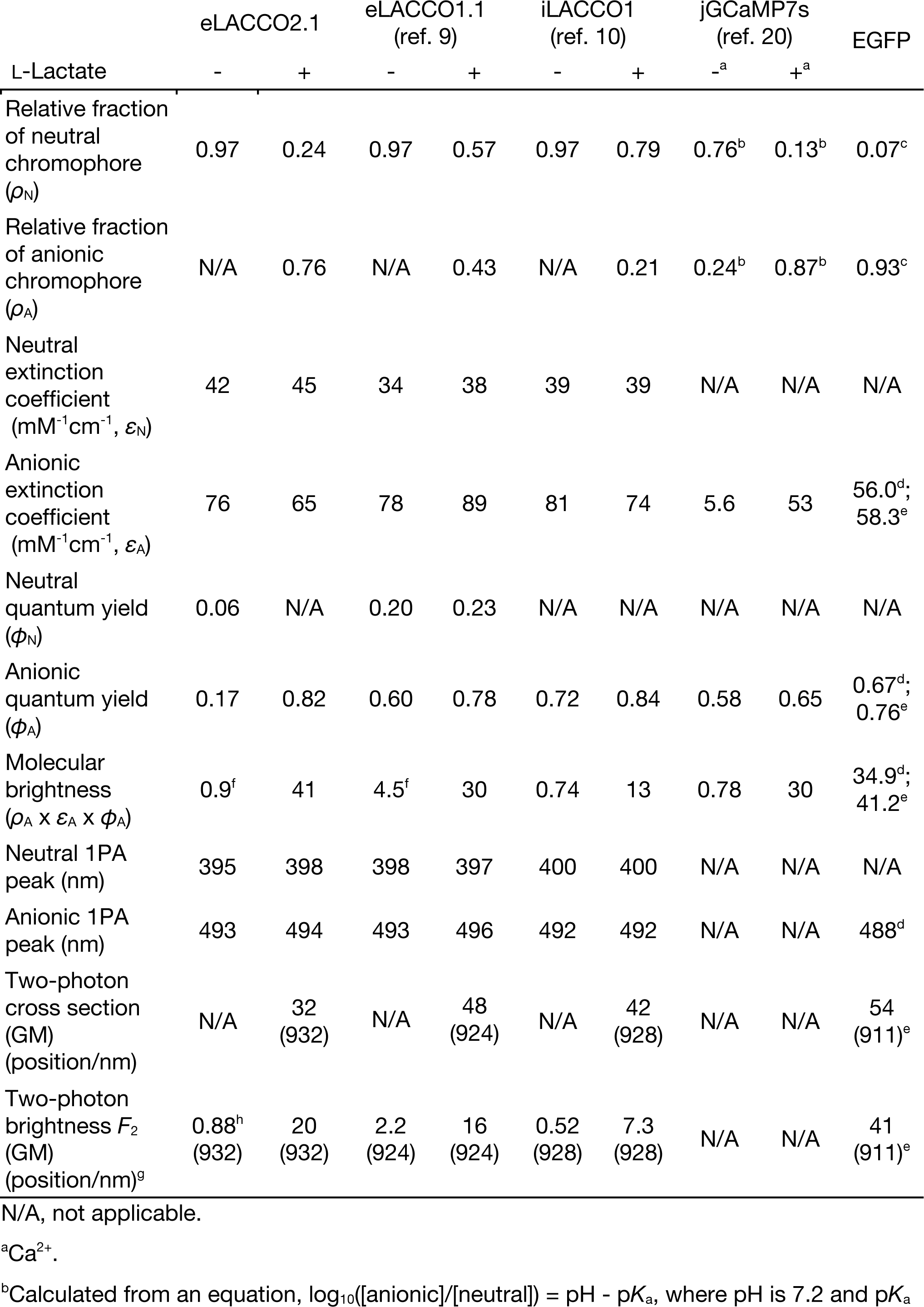
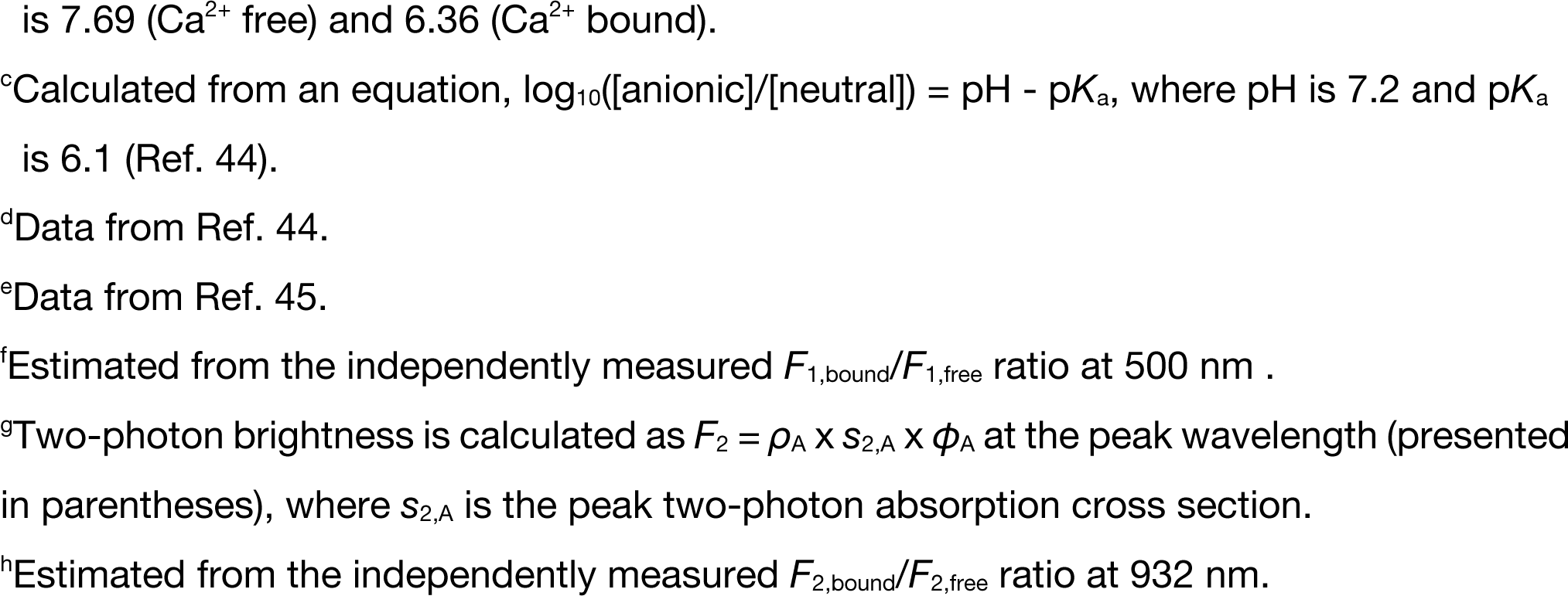
One- and two-photon photophysical parameters of eLACCO2.1.

|  | eLACCO2.1 |  | eLACCO1.1<br>(ref. 9) |  | iLACCO1<br>(ref. 10) |  | jGCaMP7s<br>(ref. 20) |  | EGFP |
| --- | --- | --- | --- | --- | --- | --- | --- | --- | --- |
| L-Lactate | - | + | - | + | - | + | -<sup>a</sup> | +<sup>a</sup> |  |
| Relative fraction of neutral chromophore ($\rho_N$) | 0.97 | 0.24 | 0.97 | 0.57 | 0.97 | 0.79 | 0.76<sup>b</sup> | 0.13<sup>b</sup> | 0.07<sup>c</sup> |
| Relative fraction of anionic chromophore ($\rho_A$) | N/A | 0.76 | N/A | 0.43 | N/A | 0.21 | 0.24<sup>b</sup> | 0.87<sup>b</sup> | 0.93<sup>c</sup> |
| Neutral extinction coefficient ($\text{mM}^{-1}\text{cm}^{-1}$, $\epsilon_N$) | 42 | 45 | 34 | 38 | 39 | 39 | N/A | N/A | N/A |
| Anionic extinction coefficient ($\text{mM}^{-1}\text{cm}^{-1}$, $\epsilon_A$) | 76 | 65 | 78 | 89 | 81 | 74 | 5.6 | 53 | 56.0<sup>d</sup>;<br>58.3<sup>e</sup> |
| Neutral quantum yield ($\phi_N$) | 0.06 | N/A | 0.20 | 0.23 | N/A | N/A | N/A | N/A | N/A |
| Anionic quantum yield ($\phi_A$) | 0.17 | 0.82 | 0.60 | 0.78 | 0.72 | 0.84 | 0.58 | 0.65 | 0.67<sup>d</sup>;<br>0.76<sup>e</sup> |
| Molecular brightness ($\rho_A \times \epsilon_A \times \phi_A$) | 0.9<sup>f</sup> | 41 | 4.5<sup>f</sup> | 30 | 0.74 | 13 | 0.78 | 30 | 34.9<sup>d</sup>;<br>41.2<sup>e</sup> |
| Neutral 1PA peak (nm) | 395 | 398 | 398 | 397 | 400 | 400 | N/A | N/A | N/A |
| Anionic 1PA peak (nm) | 493 | 494 | 493 | 496 | 492 | 492 | N/A | N/A | 488<sup>d</sup> |
| Two-photon cross section (GM) (position/nm) | N/A | 32<br>(932) | N/A | 48<br>(924) | N/A | 42<br>(928) | N/A | N/A | 54<br>(911)<sup>e</sup> |
| Two-photon brightness $F_2$ (GM) (position/nm)<sup>g</sup> | 0.88<sup>h</sup><br>(932) | 20<br>(932) | 2.2<br>(924) | 16<br>(924) | 0.52<br>(928) | 7.3<br>(928) | N/A | N/A | 41<br>(911)<sup>e</sup> |

N/A, not applicable.
<sup>a</sup>Ca<sup>2+</sup>.<sup>b</sup>Calculated from an equation, $\log_{10}([\text{anionic}]/[\text{neutral}]) = \text{pH} - \text{pK}_a$ , where pH is 7.2 and $\text{pK}_a$
is 7.69 ( $\text{Ca}^{2+}$ free) and 6.36 ( $\text{Ca}^{2+}$ bound).
<sup>c</sup>Calculated from an equation, $\log_{10}([\text{anionic}]/[\text{neutral}]) = \text{pH} - \text{p}K_a$ , where pH is 7.2 and $\text{p}K_a$ is 6.1 (Ref. 44).
<sup>d</sup>Data from Ref. 44.
<sup>e</sup>Data from Ref. 45.
<sup>f</sup>Estimated from the independently measured $F_{1,\text{bound}}/F_{1,\text{free}}$ ratio at 500 nm .
<sup>g</sup>Two-photon brightness is calculated as $F_2 = \rho_A \times s_{2,A} \times \phi_A$ at the peak wavelength (presented in parentheses), where $s_{2,A}$ is the peak two-photon absorption cross section.
<sup>h</sup>Estimated from the independently measured $F_{2,\text{bound}}/F_{2,\text{free}}$ ratio at 932 nm.

We determined that eLACCO2.1 has an apparent *K*_d_ of 960 µM and a Hill coefficient of 0.98 for l-Lactate (**Fig. 2b**). The TTHA0766 lactate binding protein contains Ca^2+^ in the l-Lactate binding pocket, rendering TTHA0766-based lactate biosensors functional only in the presence of Ca^2+^ (refs. 9,12). Indeed, we found that eLACCO2.1 has an apparent *K*_d_ of 25 μM for Ca^2+^ and exhibits practically its full response to l-Lactate at Ca^2+^ concentrations greater than 520 μM (**Fig. 2d**). eLACCO2.1 exhibits p*K*_a_ values of 6.8 and 9.0 in the presence and absence of l-Lactate, respectively (**Fig. 2e**). The control biosensor deLACCO1 showed no response to l-Lactate (**Fig. 2b**), and pH dependence that was similar to the lactate-bound state of eLACCO2.1 (**Supplementary Fig. 5b**). Investigation of the molecular specificity revealed that eLACCO2.1 is highly specific for l-Lactate over a wide array of metabolites (**Fig. 2f**). eLACCO2.1 responds to d-lactate with an apparent *K*_d_ of 8.9 mM (**Fig. 2f inset**), a concentration that is far greater than the physiological concentration (tens of μM) in plasma^21^.

### *In vitro* characterization of R-iLACCO1

We undertook a similarly detailed characterization of the biochemical and spectral properties of R-iLACCO1 and its affinity variants (**Fig. 2g–l**, **Table 2**, **Supplementary Fig. 9**, and **Supplementary Table 2**). R-iLACCO1 has absorbance peaks at 444 nm and 579 nm, indicative of the neutral (protonated) and the anionic (deprotonated) chromophore, respectively, in the absence of l-Lactate (**Fig. 2g**). As with eLACCO2.1, lactate binding results in an increase in the anionic chromophore fraction. The molecular brightness of R-iLACCO1 in the l-Lactate bound state is 94% and 40% of the R-GECO1 in Ca^2+^ bound state and the parent mApple, respectively (**Table 2**)^22, 44^. Under one-photon excitation conditions, R-iLACCO1 in the absence of l-Lactate displays an excitation peak at 574 nm and an emission peak at 604 nm. In the presence of l-Lactate, these peak positions are slightly blue-shifted to 564 nm and 594 nm, respectively (**Fig. 1h**). The absorbance peak for the anionic form undergoes a similar blue-shift from 579 nm to 561 nm (**Fig. 2g**). The two-photon excitation spectra of R-iLACCO1 show two ill-resolved peaks in both the presence and absence of l-Lactate (**Fig. 2i**), similar to those of R-GECO1 (ref. 23). R-iLACCO1 in the absence of l-Lactate displays peaks at 1080 nm and 1152 nm, while it exhibits peaks at 1048 nm and 1116 nm in the presence of l-Lactate (**Fig. 2i**). This result indicates that R-iLACCO1 blue shifts upon l-Lactate binding under two-photon excitation, consistent with behavior under one-photon excitation. The two-photon brightness of l-Lactate-bound R-iLACCO1 excited at 1048 nm, *F*_2_ = 9.2 GM, is larger than that of Ca^2+^-bound R-GECO1 (*F*_2_ = 5 GM) (**Table 2**)^23^. Similar to some red fluorescent genetically encoded biosensors^23^, the l-Lactate induced two-photon excited fluorescence change (Δ*F*_2_/*F*_2_ = 33 at 1048 nm) is larger than the one-photon excited fluorescence change (Δ*F*/*F* = 20 at 593 nm) (**Figs. 1h** and **2i**).

**Table 2.**
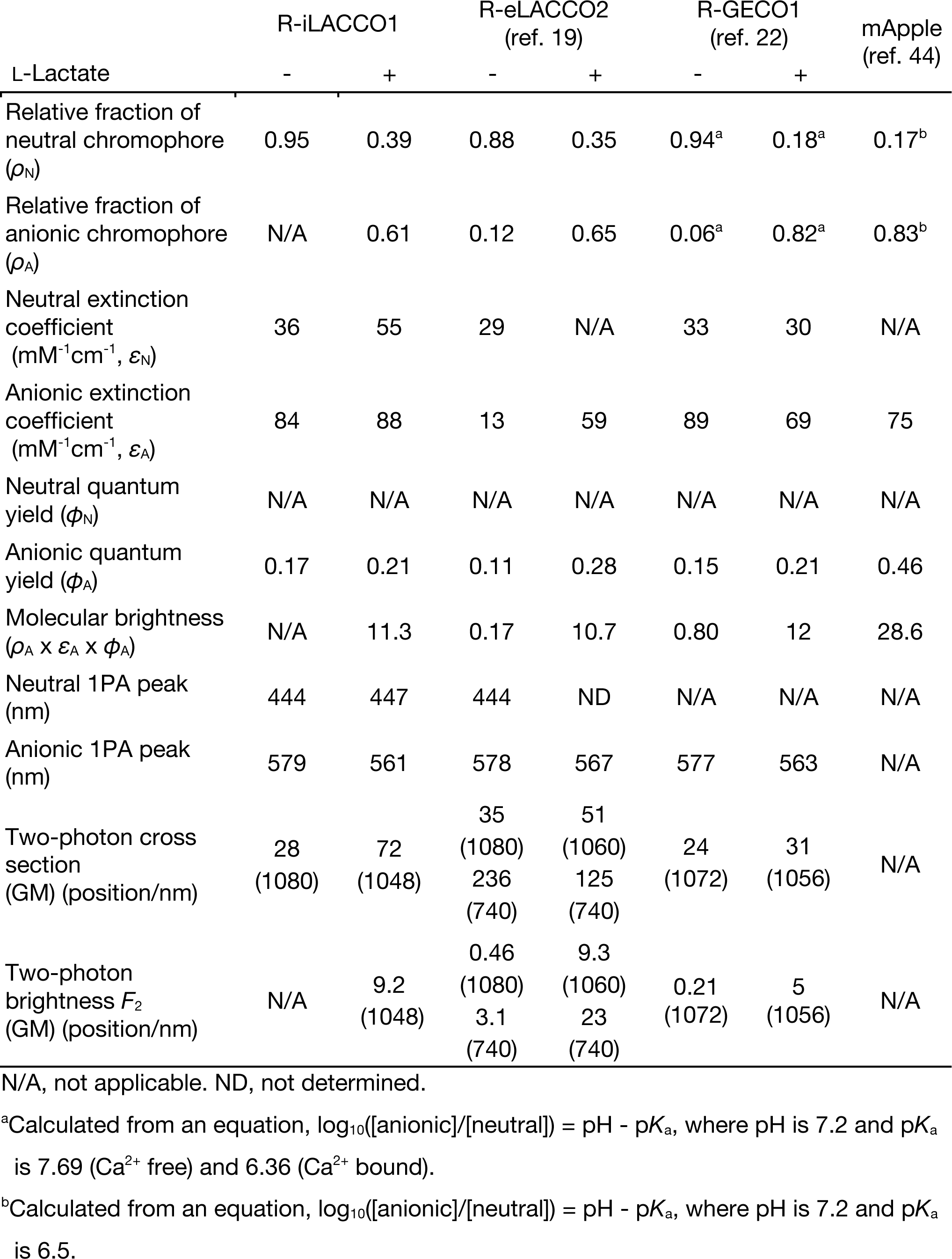
One- and two-photon photophysical parameters of R-iLACCO1.

R-iLACCO1, R-iLACCO1.1, and R-iLACCO1.2 display apparent *K*_d_s of 74 μM, 250 μM, and 350 μM for l-Lactate, respectively (**Fig. 2h**). As it incorporates the LldR l-Lactate binding domain, R-iLACCO1’s function is independent of Ca^2+^ concentration (**Fig. 2j**). R-iLACCO1 has a biphasic pH dependence (p*K*_a_ values of 5.7 and 9.6) in the presence of l-Lactate, and a monophasic pH dependence (p*K*_a_ value of 9.2) in the absence of l-Lactate (**Fig. 2k**). R-diLACCO1 showed no response to l-Lactate (**Fig. 2h**), and pH dependence that was similar to the lactate-bound state of R-iLACCO1 (**Supplementary Fig. 5d**). *In vitro* characterization revealed that R-iLACCO1 is highly specific for l-Lactate over various metabolites (**Fig. 2l**), and exhibits a substantial fluorescence response to d-lactate with an apparent *K*_d_ of 2.1 mM (**Fig. 2l inset**).

### Optimization of cell surface localization

Our previous work on eLACCO1.1 and R-eLACCO2 had demonstrated that the exact combination of N-terminal leader sequence and C-terminal anchor domain plays a critical role in determining the efficiency of cell surface targeting, as well as the functionality, of a particular extracellular biosensor^9, 19^. In parallel with the engineering of eLACCO2.1, we assessed 23 leaders^9^ and 10 anchors^24^ with eLACCO1.1 to identify the optimal combination for use in primary neurons (**Fig. 3a**). We first examined the efficiency of cell surface targeting with human CD59-derived leader sequence and 10 different anchors, including peptide-based and lipid (glycosylphosphatidylinositol, GPI)-based anchors (**Fig. 3b**). Fluorescence imaging revealed that the reticulon-4 receptor (Nogo receptor, NGR) GPI anchor resulted in the highest efficiency of cell surface localization and also exhibited a higher Δ*F*/*F* compared to the original CD59-eLACCO1.1-CD59 (**Fig. 3b,c**)^9^. We next screened various N-terminal leader sequences in combination with the NGR anchor in the presence of l-Lactate (**Fig. 3d**). Of 23 leader sequences examined, influenza hemagglutinin (HA) provided very good membrane localization and also showed greater fluorescent signal relative to CD59-eLACCO1.1-NGR in neurites (**Fig. 3e**). Having identified HA and NGR as the optimal combination for cell surface targeting, we appended these sequences to eLACCO2.1 to produce our final expression construct. For the results that follow, HA-eLACCO2.1-NGR was used for all cell imaging experiments (**Fig. 3a**).

**Fig. 3.**
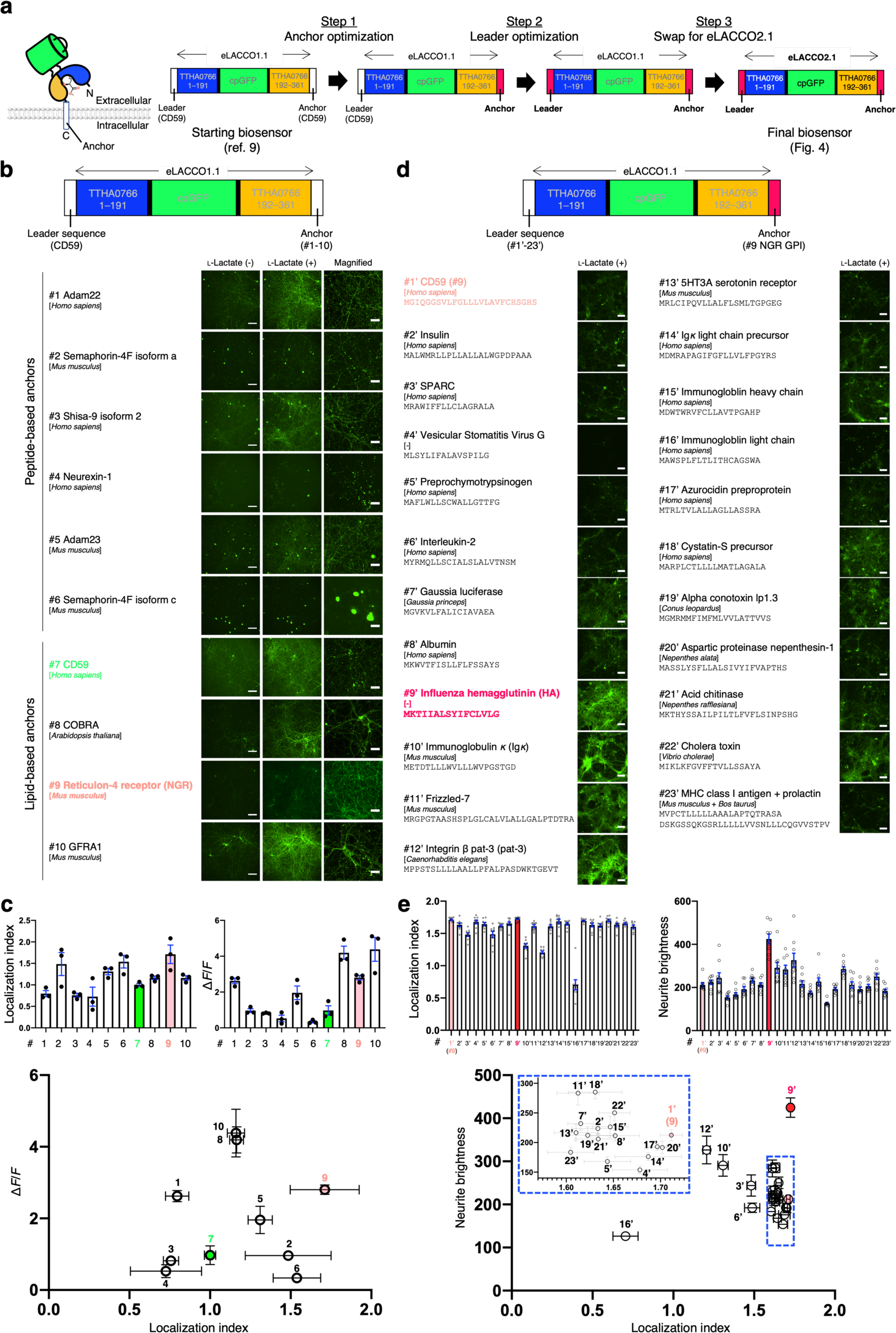
Optimization of leader/anchor combination for cell-surface expression. (**a**) Overview of the biosensor optimization. (**b**) Localization of eLACCO1.1 with CD59 leader sequence and different anchors in rat primary neurons. Scale bars, 200 μm. Right panels show magnified images in the presence of l-Lactate. Scale bars, 50 μm. (**c**) Localization index and Δ*F*/*F* of eLACCO1.1 with CD59 leader sequence and different anchors in rat primary neurons. Mean ± s.e.m., *n* = 3 field of views (FOVs) over 3 wells per construct. (**d**) Localization of eLACCO1.1 with different leader sequences and NGR GPI anchor in rat primary neurons in the presence of l-Lactate. Scale bars, 200 μm. (**e**) Localization index and neurite brightness of eLACCO1.1 with different leader sequences and NGR GPI anchor in rat primary neurons. Mean ± s.e.m., *n* = 9 FOVs over 3 wells per construct.

### Characterization of eLACCO2.1 and R-iLACCO1 in live mammalian cells

We expressed and characterized eLACCO2.1, fused with the optimized leader (HA) and anchor (NGR) sequences, in mammalian cells. eLACCO2.1 localized well on the cell membrane (**Fig. 4a**), while CD59-eLACCO1.1-CD59 formed some fluorescent puncta as previously reported^9^. The application of 10 mM l-Lactate robustly increased fluorescence intensity (Δ*F*/*F* of 8.8 ± 0.2, mean ± s.e.m.) of eLACCO2.1 expressed on HeLa cells (**Fig. 4b,c**). This response is 138% that of eLACCO1.1 (Δ*F*/*F* of 6.4 ± 0.2, mean ± s.e.m.) (**Fig. 4c**) treated using the same protocol. The control biosensor deLACCO1 had good membrane localization and, as expected, did not respond to l-Lactate even at the highest concentration tested (**Fig. 4a–d**). eLACCO2.1 has an *in situ* apparent *K*_d_ of 580 μM for l-Lactate (**Fig. 4d**). Similar to eLACCO1.1 (ref. 9), eLACCO2.1 displays Ca^2+^ dependent fluorescence with an apparent *K*_d_ of 270 μM, which is much lower than the extracellular Ca^2+^ concentration in brain tissue (1.5–1.7 mM)^25^ and serum (0.9–1.3 mM)^26^ (**Fig. 4e**). To probe the response kinetics of eLACCO2.1, we exposed eLACCO2.1-expressing HeLa cells to a solution containing 0 mM, then 10 mM, and then 0 mM l-Lactate. Imaging during the l-Lactate concentration changes revealed that eLACCO2.1 had faster on and off rates (τ_on_ of 57 ± 4 s and τ_off_ of 20 ± 2 s, mean ± s.d.) than eLACCO1.1 (τ_on_ of 74 ± 7 s and τ_off_ of 24 ± 5 s, mean ± s.d.) (**Fig. 4f**). To test photostability, we continuously illuminated eLACCO2.1-expressing HeLa cells using one-photon wide-field microscopy (∼10 mW cm^-2^, **Fig. 4g**). The photostability of eLACCO2.1 (τ_bleach_ of 189 ± 26, mean ± s.e.m.) was comparable to the parent EGFP (τ_bleach_ of 169 ± 4, mean ± s.e.m.) and higher than eLACCO1.1 (τ_bleach_ of 111 ± 6, mean ± s.e.m.) in the absence of l-Lactate, and was lower (τ_bleach_ of 55 ± 1, mean ± s.e.m.) than eLACCO1.1 (τ_bleach_ of 83 ± 7, mean ± s.e.m.) in the presence of l-Lactate. deLACCO1 was less photostable than EGFP in both the presence (τ_bleach_ of 74 ± 1, mean ± s.e.m.) and absence (τ_bleach_ of 70 ± 1, mean ± s.e.m.) of l-Lactate.

**Fig. 4.**
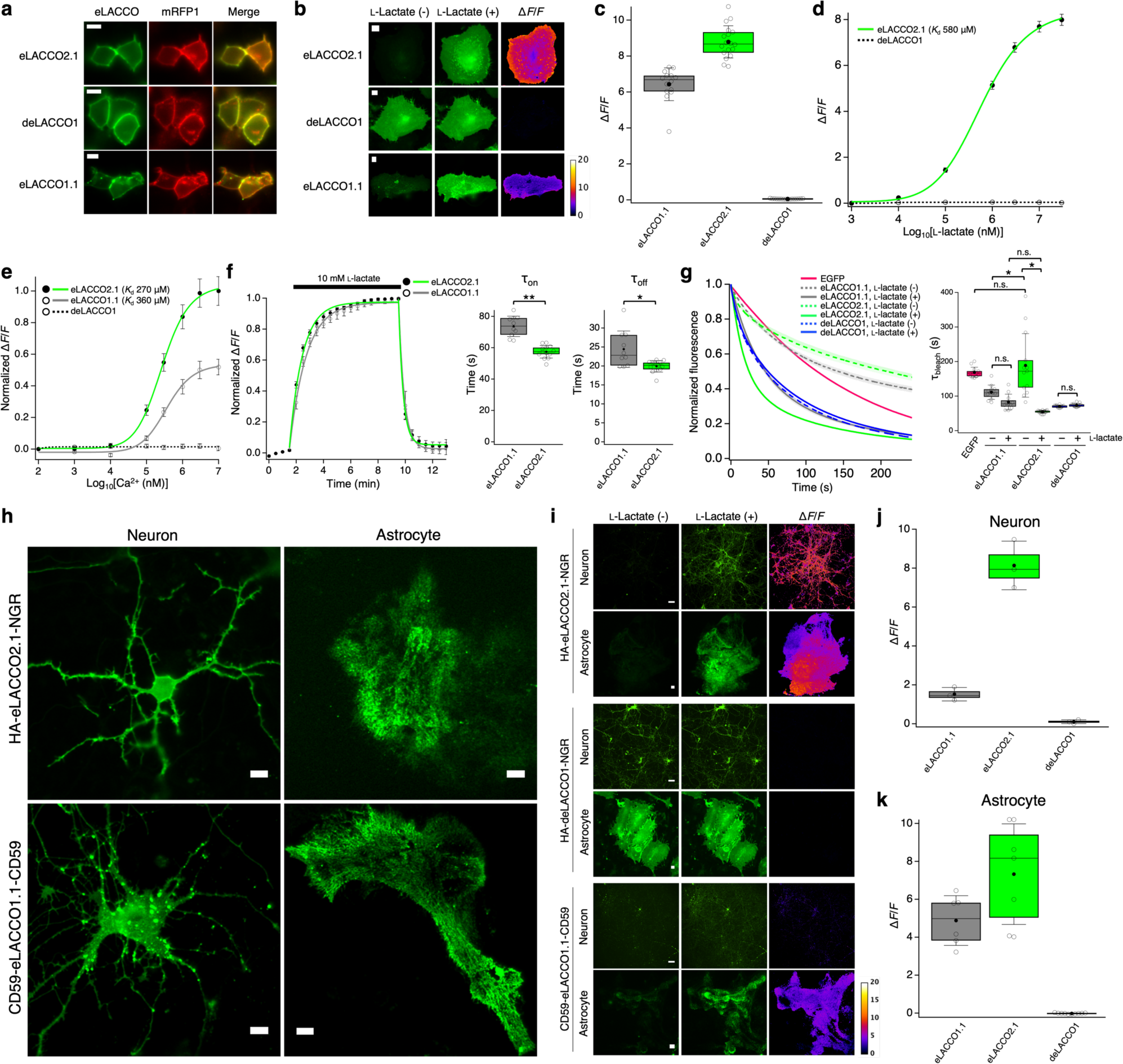
Characterization of eLACCO2.1 in live mammalian cells. (**a**) Localization of eLACCO2.1 and control biosensor deLACCO1 with the optimized leader sequence and anchor in HEK293T cells. Localization of eLACCO1.1 with CD59-derived leader and anchor domain^9^ is also shown. mRFP1 (ref. 43) with the Igκ leader sequence and PDGFR transmembrane domain was used as a cell surface marker. Scale bars, 10 μm. (**b**) Representative images of eLACCO2.1 and deLACCO1 expressed on cell surface of HeLa cells before and after 10 mM l-Lactate stimulation. Similar results were observed in more than 10 cells. Scale bars, 10 μm. (**c**) l-Lactate-dependent Δ*F*/*F* for eLACCO2.1, deLACCO1, and eLACCO1.1 expressed on HeLa cells. *n* = 18, 16, and 14 cells for eLACCO2.1, deLACCO1, and eLACCO1.1, respectively. (**d**) *In situ* l-Lactate titration of eLACCO2.1 and deLACCO1 expressed on HeLa cells. *n* = 11 and 11 cells for eLACCO2.1 and deLACCO1, respectively (mean ± s.e.m.). (**e**) *In situ* Ca^2+^ titration of eLACCO1.1, eLACCO2.1, and deLACCO1 expressed on HeLa cells in the presence (10 mM) of l-Lactate. *n* = 10, 10, and 11 cells for eLACCO1.1, eLACCO2.1, and deLACCO1, respectively (mean ± s.e.m.). (**f**) Time course of the fluorescence response of eLACCO2.1 and eLACCO1.1 on the surface of HeLa cells. The plot was fitted with a single exponential function. *n* = 13 and 10 cells for eLACCO2.1 and eLACCO1.1, respectively (mean ± s.d.). Two-tailed Student’s *t*-test. *p < 0.01 **p < 0.0001 (**g**) Photobleaching curves (left) and τ_bleach_ (right) for EGFP, eLACCO1.1, eLACCO2.1, and deLACCO1 expressed on the surface of HeLa cells using one-photon wide-field microscopy. *n* = 13, 12, 11, 13, 16, 12, and 15 cells for EGFP, eLACCO1.1 (l-Lactate -/+), eLACCO2.1 (l-Lactate -/+), and deLACCO1 (l-Lactate -/+), respectively. In the photobleaching curves, solid lines represent mean value and shaded areas represent s.e.m. One-way ANOVA followed by Tukey’s multiple comparison. *p < 0.0001. (**h**) Expression of eLACCO variants expressed on the surface of rat primary neurons and astrocytes. Scale bars, 10 μm. (**i**) l-Lactate (10 mM) response of eLACCO variants expressed on cell surface of rat primary neurons and astrocytes. Scale bars, 10 μm. (**j**) l-Lactate-dependent Δ*F*/*F* of eLACCO biosensors under hSyn promoter on rat primary neurons. *n* = 3 FOVs over 3 wells per construct. (**k**) l-Lactate-dependent Δ*F*/*F* of eLACCO biosensors under gfaABC1D promoter expressed on rat cortical astrocytes. *n* = 6, 7, and 10 cells for eLACCO2.1, deLACCO1, and eLACCO1.1, respectively. In the box plots of (c), (f), (g), (j), and (k), the horizontal line is the median; the top and bottom horizontal lines are the 25^th^ and 75^th^ percentiles for the data; and the whiskers extend one standard deviation range from the mean represented as black filled circle.

To characterize the performance of eLACCO2.1 on the surface of neurons, we expressed it under the control of human synapsin (hSyn) promoter in rat primary cortical and hippocampal neurons. We observed that neurons expressing HA-eLACCO2.1-NGR exhibited bright membrane-localized fluorescence, while the first-generation CD59-eLACCO1.1-CD59 exhibited membrane-localized fluorescence with some puncta apparent (**Fig. 4h**). Upon bath application of 10 mM l-Lactate, eLACCO2.1 had Δ*F*/*F* values of 8.1 ± 0.7 (mean ± s.e.m.), which is 5.4-fold higher than eLACCO1.1 (Δ*F*/*F* = 1.5 ± 0.2, mean ± s.e.m.) (**Fig. 4i,j**). To determine whether eLACCO2.1 could be used to detect extracellular l-Lactate concentration changes on the surface of astrocytes, we expressed it under the control of the gfaABC1D promoter in rat primary cortical and hippocampal astrocytes. Fluorescence imaging revealed that the biosensor was expressed well (**Fig. 4h**), and that eLACCO2.1 displayed improved Δ*F*/*F* values of 7.3 ± 1.0 (mean ± s.e.m., **Fig. 4i,k**) compared to eLACCO1.1 (Δ*F*/*F* = 4.9 ± 0.5, mean ± s.e.m.), upon bath application of 10 mM l-Lactate. deLACCO1 showed no response to l-Lactate, similar to our results obtained with HeLa cells and neurons. Taken together, these results indicated that eLACCO2.1 with the optimized leader and anchor has improved performance relative to eLACCO1.1 and can be used for imaging of extracellular l-Lactate concentration dynamics on the surfaces of neurons and astrocytes.

Next, we characterized R-iLACCO variants in mammalian cells. The three R-iLACCO affinity variants all robustly increased fluorescence intensity (Δ*F*/*F* of 5.5 ± 0.6, 7.6 ± 0.7, and 11 ± 0.5, for R-iLACCO1, R-iLACCO1.1, and R-iLACCO1.2, respectively, mean ± s.e.m.) upon treatment of HeLa cells with l-Lactate (**Fig. 5a, b**). Under the same conditions, the control biosensor R-diLACCO1 did not respond to l-Lactate and the Laconic FRET-based l-Lactate biosensor^11^ exhibited Δ*R*/*R* of 0.19 ± 0.02 (mean ± s.e.m.). These results confirm that R-iLACCO variants respond to l-Lactate *in situ* and that their fluorescence response to l-Lactate is much larger than Laconic. R-iLACCO1, R-iLACCO1.1, and R-iLACCO1.2 have *in situ* apparent *K*_d_s of 680 μM, 3.0 mM, and 4.0 mM for l-Lactate, respectively (**Fig. 5c**). Bath application of l-Lactate revealed that R-iLACCO1 had a faster on rate (τ_on_ of 15.8 ± 0.2 s, mean ± s.e.m.) and a slower off rate (τ_off_ of 174 ± 21 s, mean ± s.e.m.) than R-iLACCO1.1 and R-iLACCO1.2 (**Fig. 5d**).

**Fig. 5.**
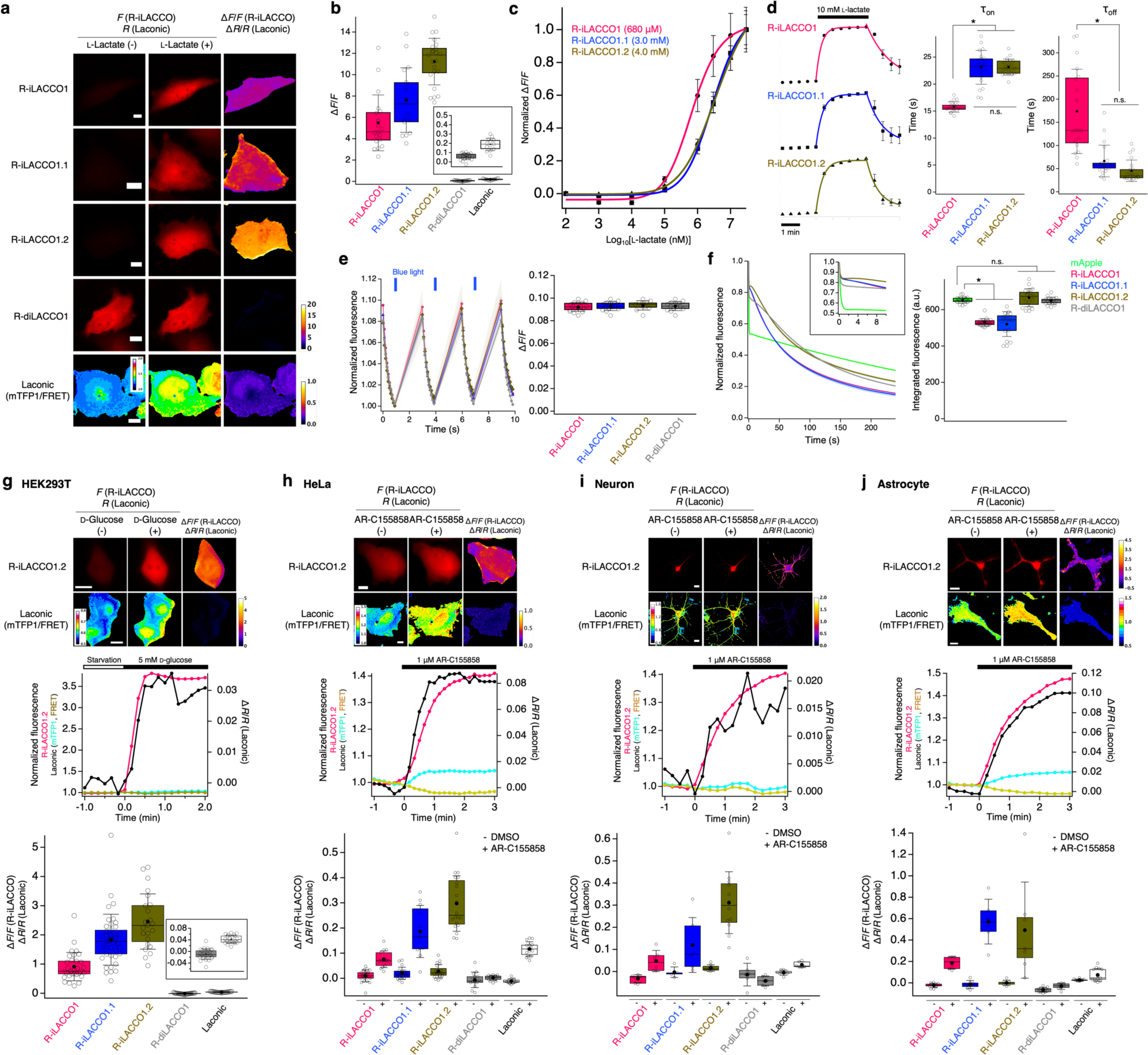
Characterization of R-iLACCO variants in live mammalian cells. (**a**) Representative images of R-iLACCO variants and Laconic expressed in HeLa cells, pretreated with 500 μM iodoacetate, 10 μM nigericine, and 2 μM rotenone to inhibit intracellular l-Lactate production and equilibrate the pH and l-Lactate, before and after 10 mM l-Lactate stimulation. Similar results were observed in more than 10 cells. Scale bars, 10 μm. (**b**) l-Lactate-dependent Δ*F*/*F* for R-iLACCO variants expressed in HeLa cells pharmacologically pretreated as shown in (a). *n* = 19, 17, 23, 20, and 13 cells for R-iLACCO1, R-iLACCO1.1, R-iLACCO1.2, R-diLACCO1, and Laconic, respectively. (**c**) *In situ* l-Lactate titration of R-iLACCO variants expressed in HeLa cells pharmacologically pretreated similar to (a). *n* = 10, 18, and 18 cells for R-iLACCO1, R-iLACCO1.1, and R-iLACCO1.2, respectively (mean ± s.e.m.). (**d**) Time course of the fluorescence response of R-iLACCO variants in HeLa cells pharmacologically pretreated similar to (a). The plot was fitted with a single exponential function. *n* = 19, 21, and 21 cells for R-iLACCO1, R-iLACCO1.1, and R-iLACCO1.2, respectively (mean ± s.d.). One-way ANOVA followed by Tukey’s multiple comparison. *p < 0.0001. (**e**) Fluorescence traces (left) and Δ*F*/*F* (right) in response to blue light illumination of HeLa cells expressing R-iLACCO variants. *n* = 17, 14, 11, and 14 cells for R-iLACCO1, R-iLACCO1.1, R-iLACCO1.2, and R-diLACCO1, respectively. Fluorescence traces represent mean ± s.e.m. (**f**) Photobleaching curves (left) and integrated fluorescence (right) for mApple and R-iLACCO variants expressed in HeLa cells using one-photon wide-field microscopy for continuous illumination. Inset shows the photobleaching curves in the first 10 seconds. *n* = 18, 20, 18, 19, and 17 cells for mApple, R-iLACCO1, R-iLACCO1.1, R-iLACCO1.2, and R-diLACCO1, respectively. In the photobleaching curves, solid lines represent mean value and shaded areas depict s.e.m. One-way ANOVA followed by Tukey’s multiple comparison. *p < 0.0001. Note that the photobleaching timescale under continuous illumination in this experiment is not applicable to that under intermittent illumination used for conventional experiments. (**g**) Representative images (top) and fluorescence traces (middle) of R-iLACCO1.1 or Laconic expressed in HEK293T upon 5 mM D-glucose treatment after starvation. Box plots (bottom) show Δ*F*/*F* for R-iLACCO variants and Δ*R*/*R* for Laconic. *n* = 31, 41, 23, 26, and 22 cells for R-iLACCO1, R-iLACCO1.1, R-iLACCO1.2, R-diLACCO1, and Laconic, respectively. Scale bars, 10 μm. (**h**) Representative images (top) and fluorescence traces (middle) of R-iLACCO1.2 or Laconic expressed in HeLa cells upon 1 μM AR-C155858 treatment. Box plots (bottom) show Δ*F*/*F* for R-iLACCO variants and Δ*R*/*R* for Laconic. *n* = 14 (22 for DMSO), 11 (19), 20 (19), 13 (14), and 15 (11) cells for R-iLACCO1, R-iLACCO1.1, R-iLACCO1.2, R-diLACCO1, and Laconic, respectively. Scale bars, 10 μm. (**i**) Representative images (top) and fluorescence traces (middle) of hSyn-R-iLACCO1.2 or CMV-Laconic expressed in primary neurons upon 1 μM AR-C155858 treatment. Box plots (bottom) show Δ*F*/*F* for R-iLACCO variants and Δ*R*/*R* for Laconic. *n* = 6 (6 for DMSO), 6 (5), 12 (8), 6 (6), and 6 (7) neurons for R-iLACCO1, R-iLACCO1.1, R-iLACCO1.2, R-diLACCO1, and Laconic, respectively. Scale bars, 20 μm. (**j**) Representative images (top) and fluorescence traces (middle) of gfaABC1D-R-iLACCO1.2 or CMV-Laconic expressed in primary astrocytes upon 1 μM AR-C155858 treatment. Box plots (bottom) show Δ*F*/*F* for R-iLACCO variants and Δ*R*/*R* for Laconic. *n* = 6 (10 for DMSO), 6 (7), 7 (7), 7 (11), and 13 (11) astrocytes for R-iLACCO1, R-iLACCO1.1, R-iLACCO1.2, R-diLACCO1, and Laconic, respectively. Scale bars, 20 μm. In the box plots of (b), (d), (e), (f), (g), (h), (i), and (j), the horizontal line is the median; the top and bottom horizontal lines are the 25^th^ and 75^th^ percentiles for the data; and the whiskers extend one standard deviation range from the mean, which is represented as a black filled circle.

Many cpmApple-based biosensors can undergo photoactivation when illuminated with blue light^27^, hampering their combined use with blue light-responsive biosensors and optogenetic tools such as GCaMP^28^ and channelrhodopsin (ChR)^29^. We therefore tested the R-iLACCO variants for photoactivation upon blue-light illumination (∼10 mW cm^-2^ at 470 nm) in HeLa cells, and found that blue-light illumination elicited a small increase (Δ*F*/*F* ∼ 0.09) in fluorescence intensity (**Fig. 5e**). This result indicates that the photoactivation and recovery of the R-iLACCO variants occurs with much faster kinetics and with far smaller Δ*F*/*F* than maximum l-Lactate dependent fluorescence response (**Fig. 5b,d**). Red fluorescent proteins (RFPs) can display complex photobleaching behavior compared to GFP^30^. To test photostability, we continuously illuminated R-iLACCO-expressing HeLa cells under wide-field microscopy (∼4 mW cm^-2^ at 560 nm, **Fig. 5f**). R-iLACCO1.2 and R-diLACCO1 exhibited photostability (integrated fluorescence, IF, of 666 ± 50 and 650 ± 20, mean ± s.d., respectively) comparable to the parent mApple RFP (IF of 653 ± 17, mean ± s.d.), whereas R-iLACCO1 and R-iLACCO1.1 exhibited lower photostability (IF of 530 ± 20 and 520 ± 68, mean ± s.d., respectively) than mApple.

To investigate whether R-iLACCO variants can enable monitoring of the dynamics of endogenous l-Lactate in mammalian cells, we performed live cell imaging of R-iLACCO variants in a variety of cell types and stimulation conditions (**Fig. 5g–j**). Glucose treatment promotes glycolysis in glucose-starved cells, leading to an increase of intracellular l-Lactate concentration^31^. Treatment of glucose-starved HEK293T cells with glucose resulted in an increase in fluorescence intensity of R-iLACCO variants (**Fig. 5g**). R-diLACCO1 showed no response in the same imaging condition, indicating that R-iLACCO variants enable monitoring of the increase in intracellular l-Lactate concentration. Notably, the low-affinity R-iLACCO1.2 showed the largest fluorescence response of all R-iLACCO variants and its Δ*F*/*F* (2.5 ± 0.2, mean ± s.e.m.) is 60 times larger than Laconic’s Δ*R*/*R* (0.042 ± 0.002, mean ± s.e.m.). Next, we aimed to examine the utility of R-iLACCO variants for detecting endogenous l-Lactate dynamics under more physiological conditions. Inhibition of monocarboxylate transporter (MCT) suppresses efflux of intracellular l-Lactate into the extracellular space, resulting in an increase in intracellular l-Lactate concentration^32^. The treatment of HeLa cells with MCT inhibitor AR-C155858 evoked robust fluorescence changes of R-iLACCO variants (Δ*F*/*F* of 0.08 ± 0.01, 0.19 ± 0.03, and 0.30 ± 0.02, for R-iLACCO1, R-iLACCO1.1, and R-iLACCO1.2, respectively, mean ± s.e.m.) and Laconic (Δ*R*/*R* of 0.12 ± 0.01, mean ± s.e.m.) (**Fig. 5h**). The control biosensor R-diLACCO1 showed no change in fluorescence intensity. To confirm the functionality of R-iLACCO variants in neural cells, we performed fluorescence imaging of primary neurons and astrocytes. Treatment of neurons and astrocytes with AR-C155858 yielded a robust increase in the fluorescence intensity of R-iLACCO variants (**Fig. 5 i,j**), similar to HeLa cells. Interestingly, the fluorescence response in astrocytes upon MCT inhibition is higher than that in HeLa cells and neurons for all R-iLACCO variants and Laconic. These imaging results suggest that the performance of lactate biosensors highly depends on cell types and stimulation conditions. Taken together, these results indicated that R-iLACCO variants could be particularly useful for imaging of intracellular l-Lactate concentration dynamics in mammalian cells.

## Discussion

Characterization of our previously reported eLACCO1.1 revealed that the biosensor harbored 57% of the neutral (dark) chromophore and 43% of the anionic (bright) chromophore in the lactate-bound state, suggesting that there could be substantial room to improve the Δ*F*/*F* by engineering increased brightness of eLACCO1.1 in the lactate-bound state (**Table 1**). Indeed, directed evolution for larger Δ*F*/*F* followed by affinity tuning led to the identification of eLACCO2.1 variant that has 24% of the neutral and 76% of the anionic chromophore and is brighter than eLACCO1.1 in the lactate bound state under both one- and two-photon excitation (**Table 1**).

When targeted to the membrane of cultured neurons using a CD59-derived leader and anchor domains (CD59-eLACCO1.1-CD59), the previously reported eLACCO1.1 exhibited fluorescence puncta that were suggestive of biosensor aggregation^9^. Previous studies suggest that the specific choice of N-terminal leader sequence and C-terminal anchor domain can play an important role in the efficiency of membrane localization of biosensors on the cell surface^9, 19, 24^. To achieve improved membrane localization with less puncta, we screened various combinations of leader and anchor domain and finally identified HA and NGR as the best leader and anchor domain, respectively. This screening revealed that the efficiency of membrane localization exhibited a stronger dependence on the anchor domains than on the leader sequence and the leader sequences seemed to have an impact on the expression level (**Fig. 3**). The best leader/anchor combination of the green fluorescent glutamate biosensor iGluSnFR3 (ref. 24) has been identified as Ig*κ* (immunoglobulin *κ* chain derived from *Mus musculus*) and NGR, respectively, based on the neuron-based screening similar to this study, whereas we have recently identified Ig*κ* and COBRA (GPI anchor derived from *Arabidopsis thaliana*) as the best leader/anchor combination for the red fluorescent extracellular l-Lactate biosensor R-eLACCO2 (ref. 19) using the screening based on HEK293 and HeLa cells. These studies suggest that the best leader/anchor combination depends on the specific biosensor and the screening method. Notably, consistent with other recent works^9, 19, 24^, the present study indicates a general propensity that lipid-based GPI anchors show better biosensor performance than peptide-based anchors such as a transmembrane domain of PDGFR (platelet-derived growth factor receptor derived from *Homo sapiens*) widely used for protein expression on cell surface. The optimized HA-eLACCO1.1-NGR expression construct has greatly improved efficiency of membrane localization, without fluorescent puncta, and expression levels compared to CD59-eLACCO1.1-CD59. These optimized properties were retained when eLACCO1.1 was switched to eLACCO2.1, with HA-eLACCO2.1-NGR substantially outperforming CD59-eLACCO1.1-CD59 in terms of membrane localization (**Fig. 4a,h**), Δ*F*/*F* (8.8 vs. 6.4 in HeLa cells, 8.1 vs. 1.5 in cultured neurons, and 7.3 vs. 4.9 in cultured astrocytes, **Fig. 4c,j,k**), and kinetics (τ_on_ of 57 s vs. 74 s, τ_off_ of 20 s vs. 24 s, **Fig.s 4f**).

Since the first report of a genetically encoded red fluorescent Ca^2+^ biosensor R-GECO1, a variety of other red biosensors have been developed^22, 33^. However, despite the remarkable progress in the development of such biosensors, the applications of genetically encoded red fluorescent biosensors have been relatively narrow compared to green fluorescent biosensors. This limitation can be attributed to the RFP-based biosensor specific properties such as smaller Δ*F*/*F*, dim fluorescence due to low molecular brightness and expression level, photoactivation upon blue-light illumination^27^, and subcellular mislocalization (e.g. accumulation in lysosomes)^34^. To overcome these difficulties, we performed extensive directed evolution to obtain variants with higher Δ*F*/*F* and brightness (**Fig. 1b**). Ultimately, we developed the R-iLACCO1 variant with Δ*F*/*F* of 20 that is one of the highest fluorescence responses for red fluorescent biosensor protein, as measured using purified protein (**Supplementary Table 2**)^19, 33^. Directed evolution with screening at 37 °C may be a contributing factor to the bright live-cell fluorescence of R-iLACCO1 (**Fig. 5**), presumably due to high molecular brightness (11.3 in lactate-bound state, **Table 2**) and efficient protein folding and chromophore maturation at 37 °C. We also confirmed that R-iLACCO1 and its affinity variants display limited amount of photoactivation upon blue-light illumination (**Fig. 5e**) and no undesirable punctate intracellular accumulation (**Fig. 5g−j**). In addition to these favorable aspects of R-iLACCO1, the availability of a variety of affinity versions provides the opportunity to probe intracellular l-Lactate dynamics over a broad range of concentrations that may differ between cell environments and types^35^.

Interestingly, l-Lactate titration revealed that R-iLACCO1 and its affinity variants have Hill coefficients of ∼1 (**Supplementary Table 2**), whereas a green fluorescent biosensor iLACCO1 based on the same LldR and cpGFP displays a Hill coefficient of 0.6 (ref. 10). These results suggest that the l-Lactate-dependent fluorescence response of iLACCO1 is negatively cooperative, but that of R-iLACCO1 variants is not cooperative. The original LldR protein exists as a dimer under physiological conditions^36^. Insertion of cpmApple into LldR may have abrogated the dimeric interaction between LldR protomers, leading to loss of cooperative interactions between the protomers. For reasons that remain unclear to us, insertion of cpGFP into the same site of LldR may have not disrupted the dimeric property in the same way that cpmApple did. Notably, the DNA binding domains of the LldR transcription factor were removed in both R-iLACCO1 and iLACCO1.

In summary, we have developed an improved variant of green fluorescent extracellular l-Lactate biosensor eLACCO2.1 and a red fluorescent intracellular l-Lactate biosensor R-iLACCO1. To date, there has been marked progress in the development of cyan-to-green hued fluorescent genetically encoded biosensors for intracellular l-Lactate^9–17, 19^. As an upgraded version of first-in-class extracellular l-Lactate biosensor eLACCO1.1, eLACCO2.1 is the latest advance in this regard. Furthermore, as the first-in-class red fluorescent biosensor for intracellular l-Lactate, R-iLACCO1 represents a major advance towards expanding the color palette for multiparameter imaging of metabolism. Overall, this update of LACCO biosensor series should provide new opportunities to investigate the emerging roles of l-Lactate in spatially and spectrally multiplexed manner.

## Methods

### General methods and materials

Synthetic DNA encoding the lactate binding bacterial protein LldR was purchased from Integrated DNA Technologies. Phusion high-fidelity DNA polymerase (Thermo Fisher Scientific) was used for routine polymerase chain reaction (PCR) amplifications, and Taq DNA polymerase (New England Biolabs) was used for error-prone PCR. Restriction endonucleases, rapid DNA ligation kits, and GeneJET miniprep kits were purchased from Thermo Fisher Scientific. PCR products and products of restriction digests were purified using agarose gel electrophoresis and the GeneJET gel extraction kit (Thermo Fisher Scientific). DNA sequences were analyzed by DNA sequence service of Fasmac Co., Ltd. Fluorescence excitation and emission spectra were recorded on a Spark plate reader (Tecan).

### Structural modeling of R-iLACCO1

The modeling structure of R-iLACCO1 was generated by AlphaFold2 (refs. 37,38) using an API hosted at the Söding lab in which the MMseqs2 server^39^ was used for multiple sequence alignment.

### Engineering of eLACCO2.1 and R-iLACCO1

A previously-reported green extracellular l-Lactate biosensor intermediate eLACCO0.9 (ref. 9) was subjected to an iterative process of library generation and screening in *E*. *coli* strain DH10B (Thermo Fisher Scientific) in LB media supplemented with 100 μg mL^-1^ ampicillin and 0.02% L-arabinose. Libraries were generated by error-prone PCR of the whole gene. Two rounds of the directed evolution led to the identification of eLACCO1.2. Cys340Ser mutation was introduced into eLACCO1.2. The resulting variant, designated eLACCO1.3, was subjected to an iterative process of library generation and screening in *E*. *coli* by error-prone PCR of the whole gene. For each round, approximately 200−400 fluorescent colonies were picked, cultured, and tested on 96-well plates under a plate reader. There were 6 rounds of screening before eLACCO2 was identified. Finally, Leu79Ile mutation was added to eLACCO2 to tune the lactate affinity. The resulting mutant was designated as eLACCO2.1. Asp444Asn mutation was added to eLACCO2 to abrogate the lactate affinity. Three rounds of directed evolution were performed to improve the brightness. The resulting control mutant was designated as deLACCO1.

The gene encoding cpmApple with N- and C-terminal linkers (DW and EREG, respectively) was amplified using a red extracellular l-Lactate biosensor R-eLACCO1 gene as template^19^, followed by insertion into the site between Leu109 and Val110 of LldR lactate binding protein in a pBAD (Invitrogen) by Gibson assembly (New England Biolabs). The resulting variant was designated as R-iLACCO0.1. N- and C-terminal linkers of R-iLACCO0.1 were deleted using Q5 high-fidelity DNA polymerase (New England Biolabs) to provide variants with different linker length. Variants were expressed in *E*. *coli*. Proteins were extracted using B-PER bacterial protein extraction reagent (Thermo Fisher Scientific) and tested for fluorescence brightness and lactate-dependent response. The most promising variant, designated as R-iLACCO0.2, was subjected to an iterative process of library generation and screening in *E*. *coli*. Libraries were generated by site-directed random mutagenesis or error-prone PCR of the whole gene. For each round, approximately 200−400 fluorescent colonies were picked, cultured, and tested on 96-well plates under a plate reader. There were 11 rounds of screening before R-iLACCO1, R-iLACCO1.1, and R-iLACCO1.2 were identified. Asp69Asn mutation was added to R-iLACCO1 to abrogate the lactate affinity. Two rounds of directed evolution were performed to improve the brightness. The resulting control mutant was designated as R-diLACCO1.

### Protein purification and *in vitro* characterization

The gene encoding eLACCO2.1 and R-iLACCO variants, with a poly-histidine tag on the N-terminus, was expressed from the pBAD vector. Bacteria were lysed with a cell disruptor (Branson) and then centrifuged at 15,000*g* for 20 min, and proteins were purified by Ni-NTA affinity chromatography (Agarose Bead Technologies). Absorption spectra of the samples were collected with a Lambda950 Spectrophotometer (PerkinElmer). To perform pH titrations for eLACCO2.1, protein solutions were diluted into buffers (pH from 4 to 11) containing 30 mM trisodium citrate, 30 mM sodium borate, 30 mM MOPS, 100 mM KCl, 1 mM CaCl_2_, and either no l-Lactate or 100 mM l-Lactate. To perform pH titrations for R-iLACCO variants, protein solutions were diluted into buffers (pH from 4 to 11) containing 30 mM trisodium citrate, 30 mM sodium borate, 30 mM MOPS, 100 mM KCl, and either no l-Lactate or 100 mM l-Lactate. Fluorescence intensities as a function of pH were then fitted by a sigmoidal binding function to determine the p*K*_a_. For lactate titration of eLACCO2.1, buffers were prepared by mixing an l-Lactate (-) buffer (30 mM MOPS, 100 mM KCl, 1 mM CaCl_2_, pH 7.2) and an l-Lactate (+) buffer (30 mM MOPS, 100 mM KCl, 1 mM CaCl_2_, 100 mM l-Lactate, pH 7.2) to provide l-Lactate concentrations ranging from 0 mM to 100 mM at 25 °C. For lactate titration of R-iLACCO variants, buffers were prepared by mixing an l-Lactate (-, w/o Ca^2+^) buffer (30 mM MOPS, 100 mM KCl, pH 7.2) and an l-Lactate (+, w/o Ca^2+^) buffer (30 mM MOPS, 100 mM KCl, 100 mM l-Lactate, pH 7.2) to provide l-Lactate concentrations ranging from 0 mM to 100 mM at 25 °C. Fluorescence intensities were plotted against l-Lactate concentrations and fitted by a sigmoidal binding function to determine the Hill coefficient and apparent *K*_d_. For Ca^2+^ titration, buffers were prepared by mixing a Ca^2+^ (-) buffer (30 mM MOPS, 100 mM KCl, 10 mM EGTA, 100 mM l-Lactate, pH 7.2) and a Ca^2+^ (+) buffer (30 mM MOPS, 100 mM KCl, 10 mM CaEGTA, 100 mM l-Lactate, pH 7.2) to provide Ca^2+^ concentrations ranging from 0 μM to 39 μM at 25 °C. Buffers with Ca^2+^ concentrations more than 39 μM was prepared by mixing a Ca^2+^ (-, w/o EGTA) buffer (30 mM MOPS, 100 mM KCl, 100 mM l-Lactate, pH 7.2) and a Ca^2+^ (+, w/o EGTA) buffer (30 mM MOPS, 100 mM KCl, 100 mM CaCl_2_, 100 mM l-Lactate, pH 7.2).

Two-photon excitation spectra and two-photon absorption cross sections of eLACCO2.1 were measured as described previously^9^. For R-iLACCO1, we measured the two-photon excitation spectra and two-photon absorption cross sections using our standard procedures described in ref. 40. Briefly, a tunable femtosecond laser InSight DeepSee (Spectra-Physics, Santa Clara, CA) was used to excite fluorescence of the sample contained within a PC1 Spectrofluorometer (ISS, Champaign, IL). To measure the two-photon excitation spectral shapes, we used short-pass filters 633SP and 770SP in the emission channel. LDS 798 in 1:2 CHCl3:CDCl3 and Coumarin 540A in DMSO were used as spectral standards. Quadratic power dependence of fluorescence intensity in the proteins and standards was checked at several wavelengths across the spectrum. The two-photon cross section (σ_2_) of the anionic form of the chromophore was measured as described previously^41^. Rhodamine 6G in MeOH was used as a reference standard with excitation at 1060 nm (ref. 40). For one-photon excitation we use a 532-nm line of diode laser (Coherent). A combination of filters 770SP and 561LP was used in the fluorescence channel for these measurements. Extinction coefficients were determined by alkaline denaturation as detailed in ref. 23. The two-photon absorption spectra were normalized to the measured σ_2_ values. To normalize to the total two-photon brightness (*F*_2_), the spectra were then multiplied by the quantum yield and the relative fraction of the respective form of the chromophore for which the σ_2_ was measured. The data is presented this way because R-iLACCO1 contains a mixture of the neutral and anionic forms of the chromophore. The method is described in further detail in refs. 23,42.

### Construction of mammalian expression vectors

For cell surface expression, the genes encoding eLACCO2.1 were amplified by PCR followed by digestion with BglII and EcoRI, and then ligated into a pcDNA3.1 vector (Thermo Fisher Scientific) that contains N-terminal leader sequence and C-terminal anchor. N-terminal leader sequences were derived from ref. 9 and C-terminal anchors stemmed from refs. 9 and 24. To construct R-iLACCO variants for intracellular expression, the gene encoding R-iLACCO variants in the pBAD vector was digested with XhoI and HindIII, and then ligated into pcDNA3.1 vector. To construct plasmids for expression in primary neurons, astrocytes, *ex vivo*, and *in vivo*, the gene encoding R-iLACCO variants or eLACCO variants including the leader and anchor sequence in the pcDNA3.1 vector was first amplified by PCR followed by digestion with NheI and XhoI for eLACCO and BamHI and HindIII for R-iLACCO, and then ligated into a pAAV plasmid containing the human synapsin (hSyn) or gfaABC1D promoter.

### Imaging of eLACCO and R-iLACCO variants in HeLa and HEK293T cell lines

HeLa and HEK293T cells were maintained in Dulbecco’s modified Eagle medium (DMEM; Nakalai Tesque) supplemented with 10% fetal bovine serum (FBS; Sigma-Aldrich) and 1% penicillin-streptomycin (Nakalai Tesque) at 37 °C and 5% CO_2_. Cells were seeded in 35-mm glass-bottom cell-culture dishes (Iwaki) and transiently transfected with the constructed plasmid using polyethyleneimine (Polysciences). Transfected cells were imaged using a IX83 wide-field fluorescence microscopy (Olympus) equipped with a pE-300 LED light source (CoolLED), a 40x objective lens (numerical aperture (NA) = 1.3; oil), an ImagEM X2 EM-CCD camera (Hamamatsu), Cellsens software (Olympus), and an STR stage incubator (Tokai Hit). The filter sets used in live cell imaging had the following specification. eLACCO variants and EGFP: excitation 470/20 nm, dichroic mirror 490-nm dclp, and emission 518/45 nm; R-iLACCO variants, mRFP1, and mApple: excitation 545/20 nm, dichroic mirror 565-nm dclp, and emission 598/55 nm; Laconic (CFP): excitation 438/24 nm, dichroic mirror 458-nm dclp, and emission 483/32 nm; Laconic (FRET): excitation 438/24 nm, dichroic mirror 458-nm dclp, and emission 542/27 nm. Fluorescence images were analyzed with ImageJ software (National Institutes of Health).

For photostability test, HeLa cells transfected with respective genes were illuminated by excitation light at 100% intensity of LED (∼10 mW cm^-2^ and ∼4 mW cm^-2^ on the objective lens for eLACCO and R-iLACCO variants, respectively) and their fluorescence images were recorded for 4 min with the exposure time of 50 ms and no interval time.

For imaging of lactate-dependent fluorescence, HeLa cells seeded onto coverslips were transfected with eLACCO or R-iLACCO variants. Forty-eight hours after transfection, the coverslips were transferred into Attofluor™ Cell Chamber (Thermo Fisher Scientific) with Hank’s balanced salt solution (HBSS(+); Nakalai Tesque) supplemented with 10 mM HEPES and 1 μM AR-C155858 (Tocris). Exchange of bath solutions during the image was performed in a remove-and-add manner using a homemade solution remover^9^.

For imaging of Ca^2+^-dependent fluorescence, HeLa cells on coverslips were transferred into Attofluor^TM^ Cell Chamber with HBSS(-) buffer (Nakalai Tesque) supplemented with 10 mM HEPES and 10 mM l-Lactate, 48 hours after transfection of eLACCO variants. Other bath solutions were supplemented with Ca^2+^ of 100 nM, 1 μM, 10 μM, 100 μM, 300 μM, 1 mM, 3 mM, and 10 mM. Exchange of bath solutions during the image was performed in the same way as the imaging of lactate-dependent fluorescence.

### Imaging of eLACCO and R-iLACCO variants in primary neurons and astrocytes

The neuron imaging was previously described^9^. In short, rat cortical/hippocampal primary cultures from the P0 pups (pooled tissues from males and females) were plated in glass-bottom 24-well plates where 0.5 × 10^6^ cells were used for three wells. Cultures were nucleofected at time of plating, and imaged 14 days later. For the imaging of primary astrocytes, male and female pups were obtained from a single timed-pregnant Sprague Dawley rat (Charles River Laboratories, purchased from Japan SLC, Inc.). Experiments were performed with cortical/hippocampal primary cultures from the E21 (after C-section of the pregnant rat) plated in glass-bottom 24-well plates (Cellvis) where 0.5 × 10^6^ cells were used for one well. Cultures were nucleofected at time of plating with Nucleofector 4D (Lonza), and imaged 5 days later. Astrocytes were cultured in DMEM supplemented with 10% FBS and 1% penicillin-streptomycin at 37 °C and 5% CO_2_. Culture media were replaced with 1 mL of imaging buffer (145 mM NaCl, 2.5 mM KCl, 10 mM D-glucose, 10 mM HEPES, 2 mM CaCl_2_, 1 mM MgCl_2_, pH 7.4) for imaging.

### Packaging and purification of adeno-associated viruses (AAVs)

AAVs were generated in HEK293T17 cells by a triple transfection of a helper plasmid pXX680, a plasmid encoding either the AAV2/5 or AAV2/9 rep/cap, and a pAAV plasmid encoding the corresponding LACCO biosensor. Forty-eight hours post transfection, the cells were harvested by gentle scraping and pelleted by centrifugation. Viral particles were released by 4 freeze-thaw cycles on dry-ice/ethanol and free DNA was digested with benzonase. The AAV particles were purified by a discontinuous gradient of iodixanol (15-25-40-60%) and ultracentrifugation (67,000 rpm for 70 min at 16 °C). The viral preparation was then washed and concentrated through a 100 kD amicon filter. During the last concentration step, the buffer was adjusted to 5% D-sorbitol and 0.001% pluronic acid for storage and experimentation. Titration was performed by TaqMan digital droplet-PCR using primers specific for AAV2 inverted terminal repeat (ITR).

### Animal care

For experiments performed at The University of Tokyo, all methods for animal care and use were approved by the institutional review committees of School of Science, The University of Tokyo. For experiments at HHMI Janelia Research Campus, all surgical and experimental procedures were in accordance with protocols approved by the HHMI Janelia Research Campus Institutional Animal Care and Use Committee and Institutional Biosafety Committee.

### Statistics and reproducibility

All data are expressed as mean ± s.d. or mean ± s.e.m., as specified in figure legends. Box plots are used for Figs. 4c, f, g, j, k, and 5 b, d, e, f, g, h, i, j. In these box plots, the horizontal line is the median; the top and bottom horizontal lines are the 25th and 75th percentiles for the data; and the whiskers extend one standard deviation range from the mean represented as black filled circle. Sample sizes (n) are listed with each experiment. No samples were excluded from analysis and all experiments were reproducible. Statistical analysis was performed using two-tailed Student’s *t*-test or one-way analysis of variance (ANOVA) with the Dunnett’s post hoc tests (GraphPad Prism 9), as specified in figure legends. Microsoft Excel software was used to plot for Figs. 1d,e,g,h and 2a,c,f,g,i,l.

## Data availability

The plasmids and AAVs encoding eLACCO2 and R-iLACCO1 variants that support the findings of this study are available from the corresponding authors on reasonable request.

## Acknowledgments

The authors thank S. Hario and T. Terai for technical support. Work at the University of Tokyo was supported by the Japan Society for the Promotion of Science (Grants-in-Aid for Early-Career Scientists 19K15691 and 21K14738, and Grants-in-Aid for Scientific Research S 19H05633), the Japan Science and Technology Agency (PRESTO, JPMJPR22E9), Toyota Physical and Chemical Research Institute, and The Precise Measurement Technology Promotion Foundation. Work at Montana State University was supported by National Institutes of Health (NIH) grants U01 NS094246, U24 NS109107, and F31 NS108593.

## Author Contributions

Y.N. developed eLACCO2.1 and performed *in vitro* characterization. Y.N. and G.L. developed R-iLACCO variants. Y.N. performed *in vitro* characterization of R-iLACCO variants. Y.N., Y.K., A.A., and K.P. performed screening of leader and anchor sequences in primary neurons. M.D. measured one-photon absorbance spectra and two-photon excitation spectra. M.B. and M.-E.P produced AAVs. Y.N. and R.E.C. supervised research. Y.N. and R.E.C. wrote the manuscript.

## Supplementary Figures

**Supplementary Figure 1.**
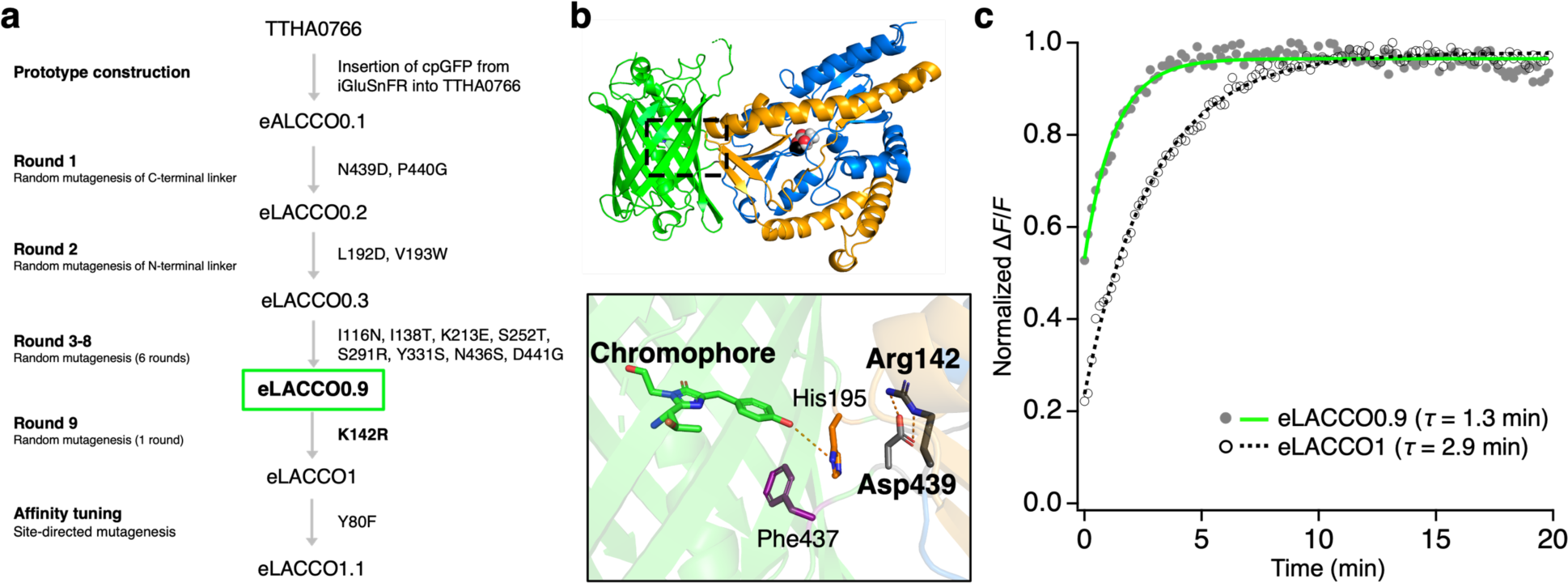
eLACCO0.9 exhibits a faster response than eLACCO1. (**a**) Lineage of eLACCO1 variants (ref. 1). Final round (round 9) of directed evolution via random mutagenesis added Lys142Arg mutation into eLACCO0.9 to produce eLACCO1. (**b**) Crystal structure (PDE 7E9Y) of eLACCO1 in l-Lactate bound state. Inset represents a zoom-in view around the chromophore. Arg142 forms a salt bridge with Asp439. (**c**) Time courses of fluorescence response of eLACCO0.9 and eLACCO1. l-Lactate (final concentration of 10 mM) was rapidly mixed at t = 0 (dead time ∼ 10 s) with the crude protein extracts. Curve fitting was performed using Δ*F*/*F* = A + B ・ e^-t/*τ*^, where *τ* is a time constant.

**Supplementary Figure 2.**
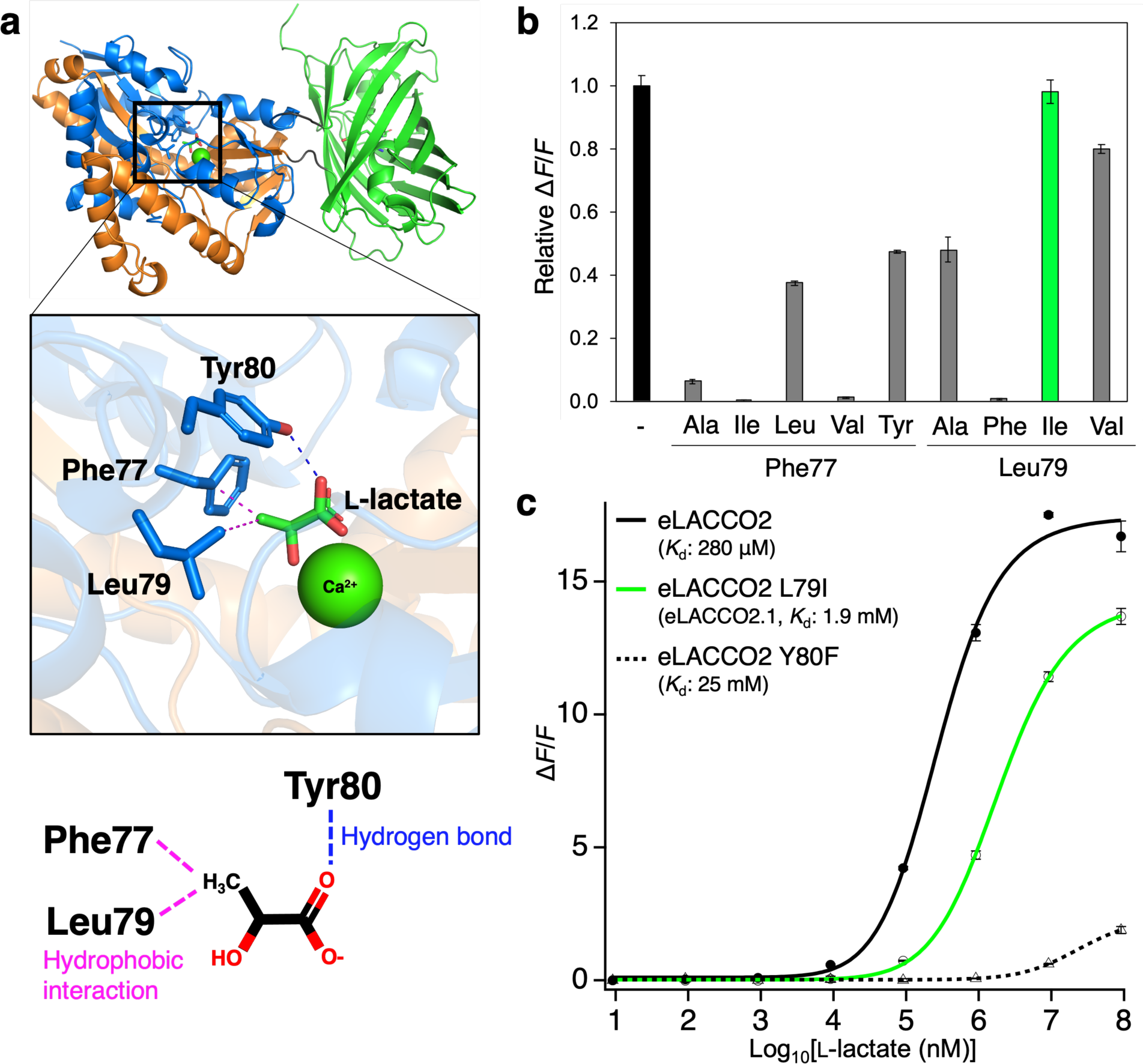
Development of affinity-tuned eLACCO2.1. (**a**) Crystal structure of eLACCO1 (PDB 7E9Y)^1^ and zoom-in view of the l-Lactate binding pocket. The phenol group of the Tyr80 side chain forms a hydrogen bond to the carboxylate group of l-Lactate. Hydrophobic side chains of Phe77 and Leu79 interact with a methyl group of l-Lactate. (**b**) Relative Δ*F*/*F* of a range of eLACCO2 variants. *n* = 3 experimental triplicates (mean ± s.d.). (**c**) Dose-response curves of eLACCO2, eLACCO2 Leu79Ile (eLACCO2.1), and eLACCO2 Tyr80Phe for l-Lactate. *n* = 3 experimental triplicates (mean ± s.d.).

**Supplementary Figure 3.**
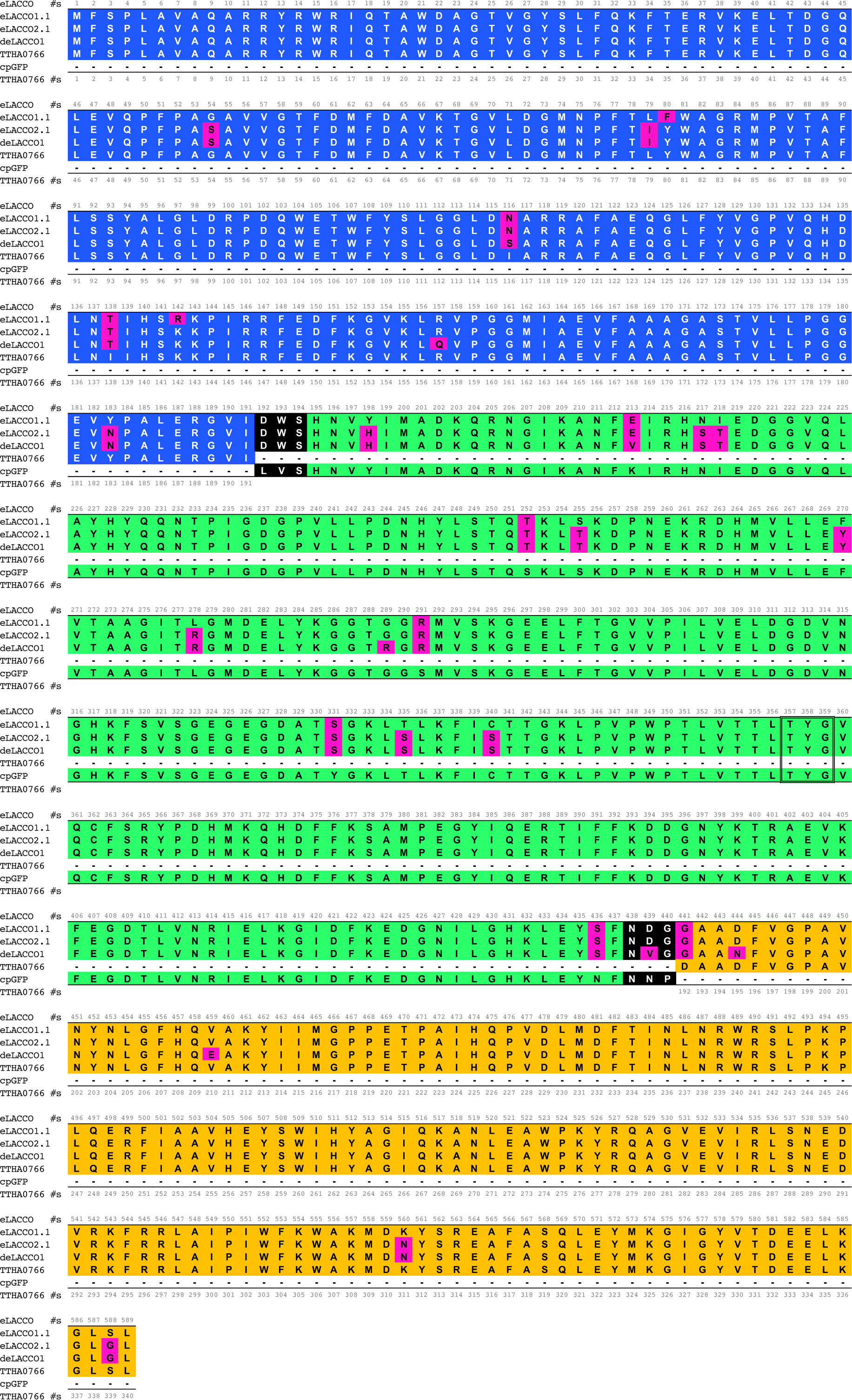
Sequence alignment of TTHA0766, cpGFP, eLACCO1.1, eLACCO2.1, and deLACCO1. Mutations in eLACCO variants, relative to TTHA0766 and cpGFP, are highlighted in magenta. The chromophore-forming residues are surrounded by a double line.

**Supplementary Figure 4.**
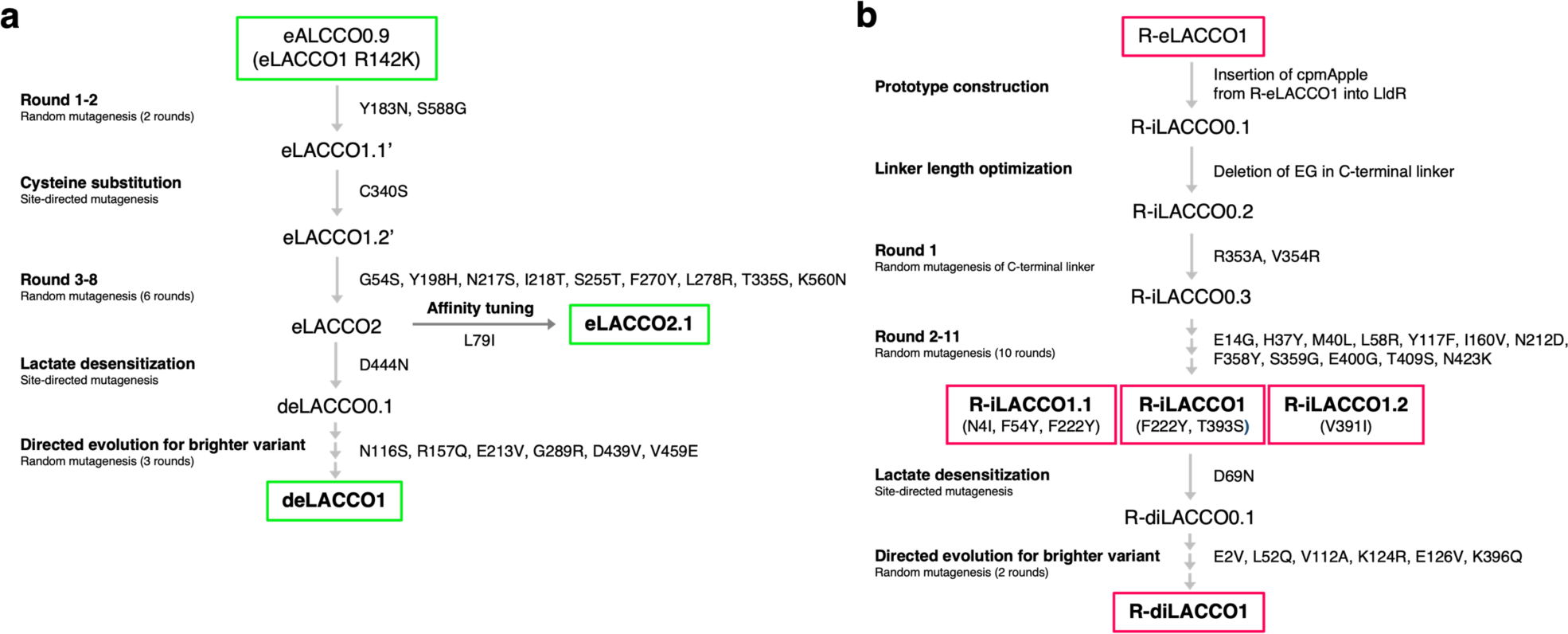
Lineage of eLACCO and R-iLACCO variants.

**Supplementary Figure 5.**
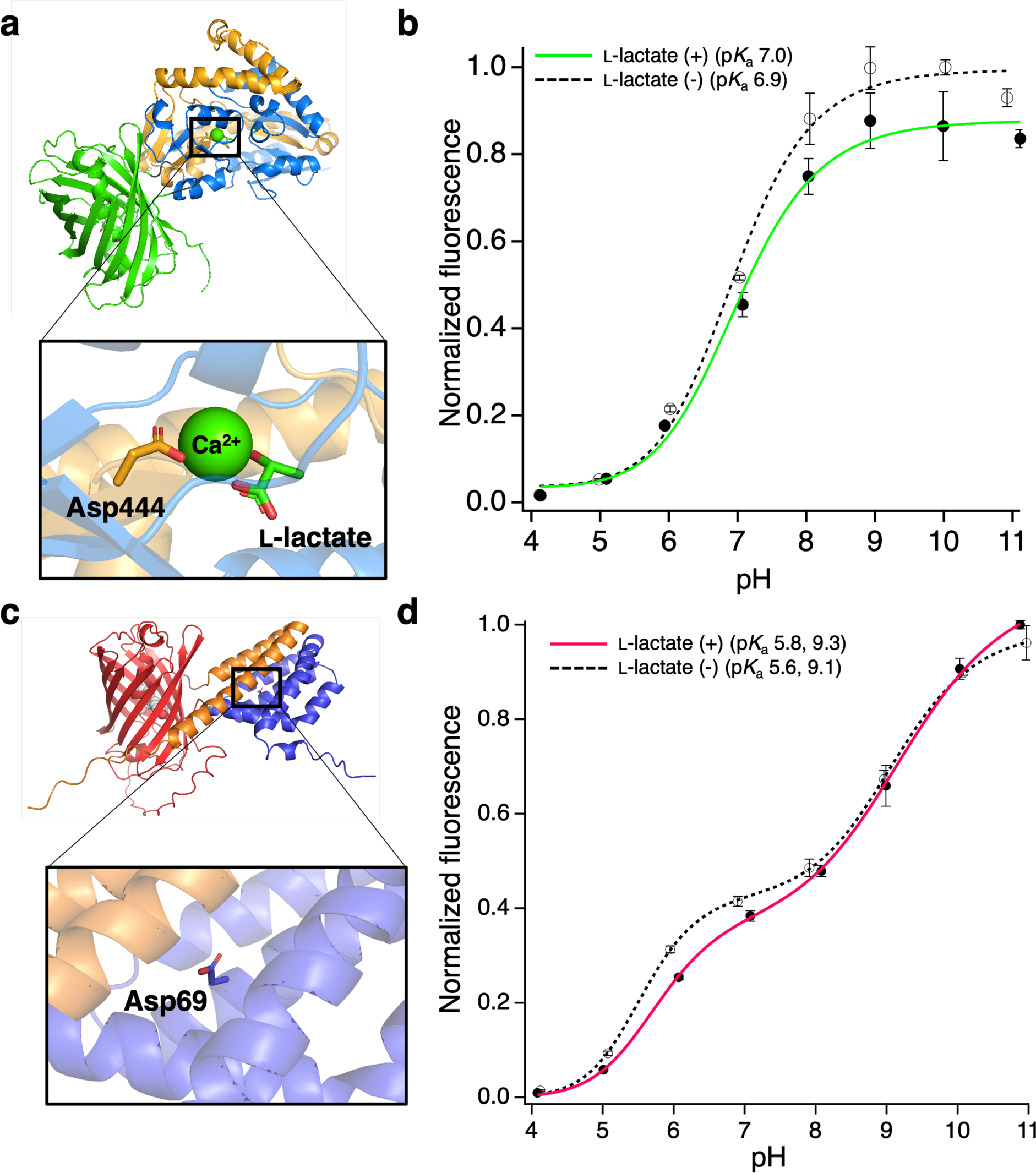
*In vitro* characterization of deLACCO1 and R-diLACCO1. (**a**) Crystal structure of eLACCO1 (ref. 1) and zoom-in view of the Ca^2+^ binding pocket. Carboxyl group of Asp444 side chain coordinates to Ca^2+^. (**b**) pH titration curves of purified deLACCO1 in the presence (100 mM) and absence of l-Lactate. n = 3 experimental triplicates (mean ± s.d.). (**c**) AlphaFold model of R-iLACCO1 and zoom-in view of the putative l-Lactate binding pocket. (**d**) pH titration curves of purified R-diLACCO1 in the presence (100 mM) and absence of l-Lactate. n = 3 experimental triplicates (mean ± s.d.).

**Supplementary Figure 6.**
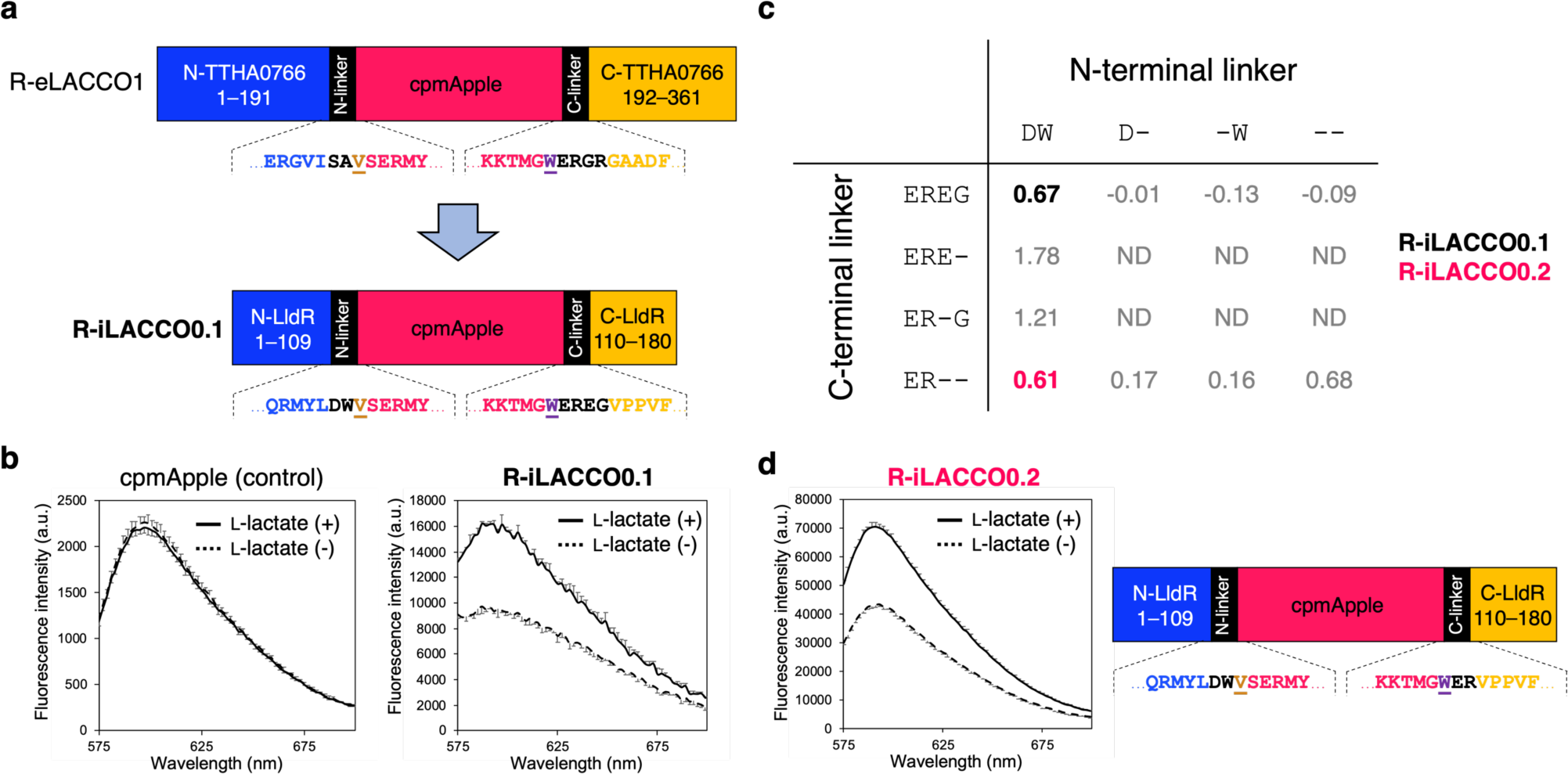
Construction of the R-iLACCO prototype. (**a**) Schematic illustration of the biosensor prototype construction based on R-eLACCO1 (ref. 2). Linker regions are shown in black and the two “gate post” residues^3^ in cpmApple are highlighted in dark orange and purple. (**b**) Emission spectra of R-iLACCO0.1 and cpmApple in the presence (10 mM) and absence of l-Lactate. Error bars represent standard deviation of triplicates. (**c**) Summary of Δ*F*/*F* of R-iLACCO variants with different linker length. ND, not determined. (**d**) Emission spectra of R-iLACCO0.2 in the presence (10 mM) and absence of l-Lactate. Right figure shows a schematic representation of the primary structure of R-iLACCO0.2. Error bars represent standard deviation of triplicates.

**Supplementary Figure 7.**
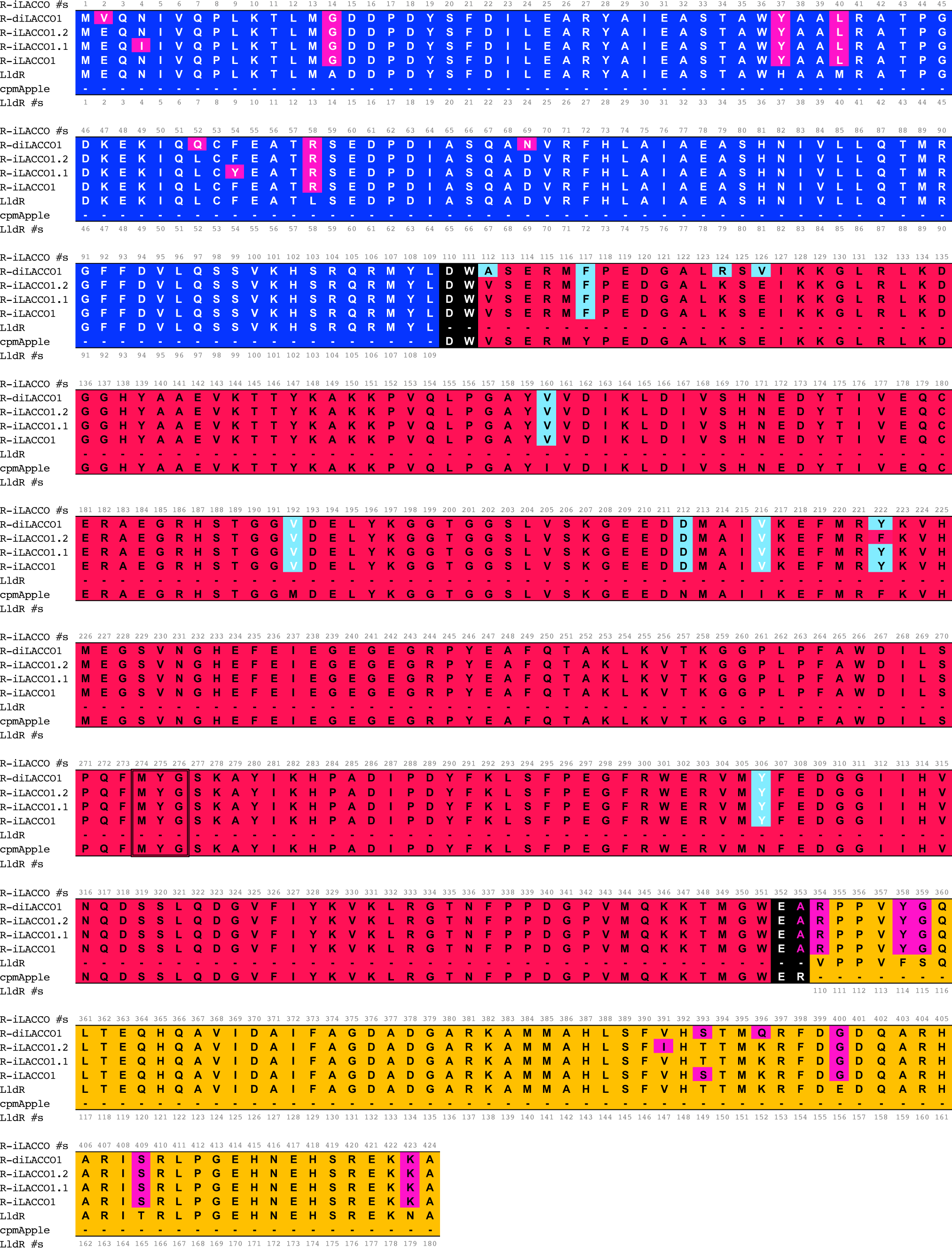
Sequence alignment of LldR, cpmApple, R-iLACCO1, R-iLACCO1.1, R-iLACCO1.2, and R-diLACCO1. Mutations in R-iLACCO variants, relative to LldR and cpmApple, are highlighted in magenta and light blue, respectively. White residues in light blue represent mutations in cpmApple derived from R-eLACCO1 (ref. 2). The chromophore-forming residues are surrounded by a double line.

**Supplementary Figure 8.**
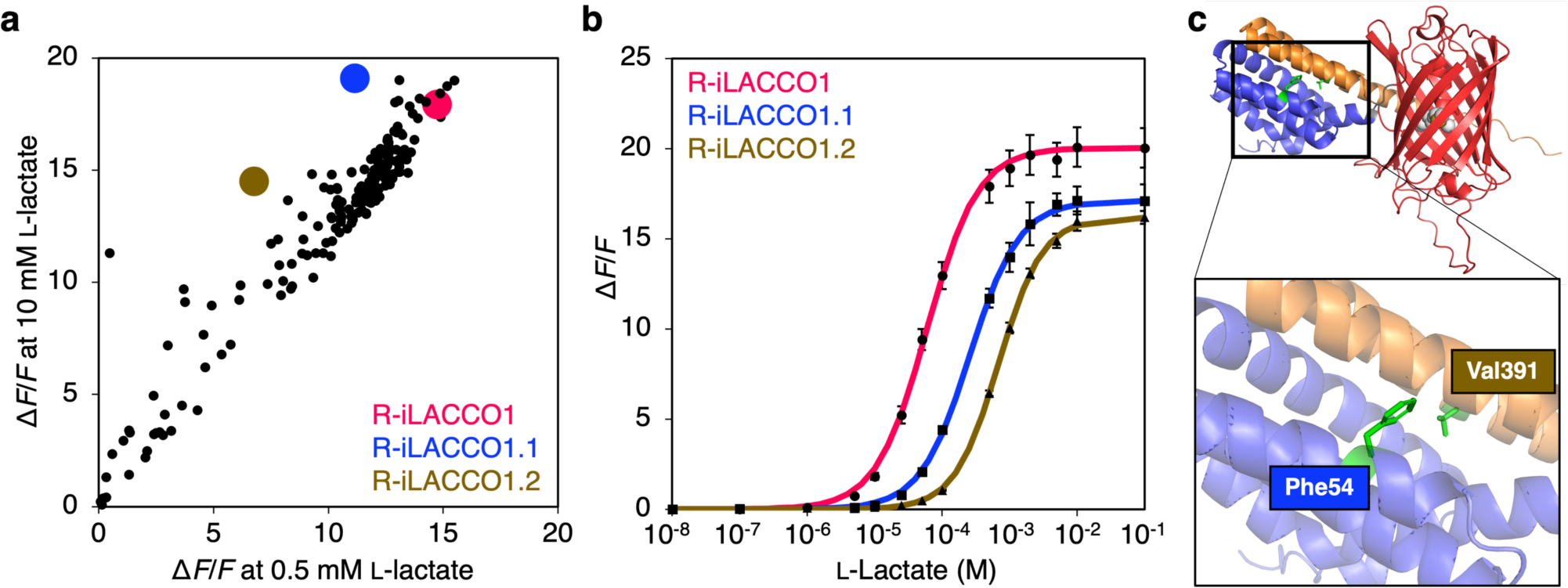
Development of R-iLACCO variants with lower l-Lactate affinity. (**a**) Scatter plot representing Δ*F*/*F* of all proteins tested in the final round of directed evolution for R-iLACCO. Δ*F*/*F* for each protein was measured at both 0.5 mM and 10 mM l-Lactate. Red, blue, and brown dot indicate R-iLACCO1, R-iLACCO1.1, and R-iLACCO1.2, respectively. (**b**) Dose-response curves of R-iLACCO1, R-iLACCO1.1 and R-iLACCO1.2 for l-Lactate. *n* = 3 experimental triplicates (mean ± s.d.). (**c**) AlphaFold model of R-iLACCO1 and zoom-in view of the putative l-Lactate binding pocket. Note that, relative to R-iLACCO1, the low affinity variants R-iLACCO1.1 and R-iLACCO1.2 have F54Y and V391I, respectively, both of which are toward the putative l-Lactate binding pocket.

**Supplementary Figure 9.**
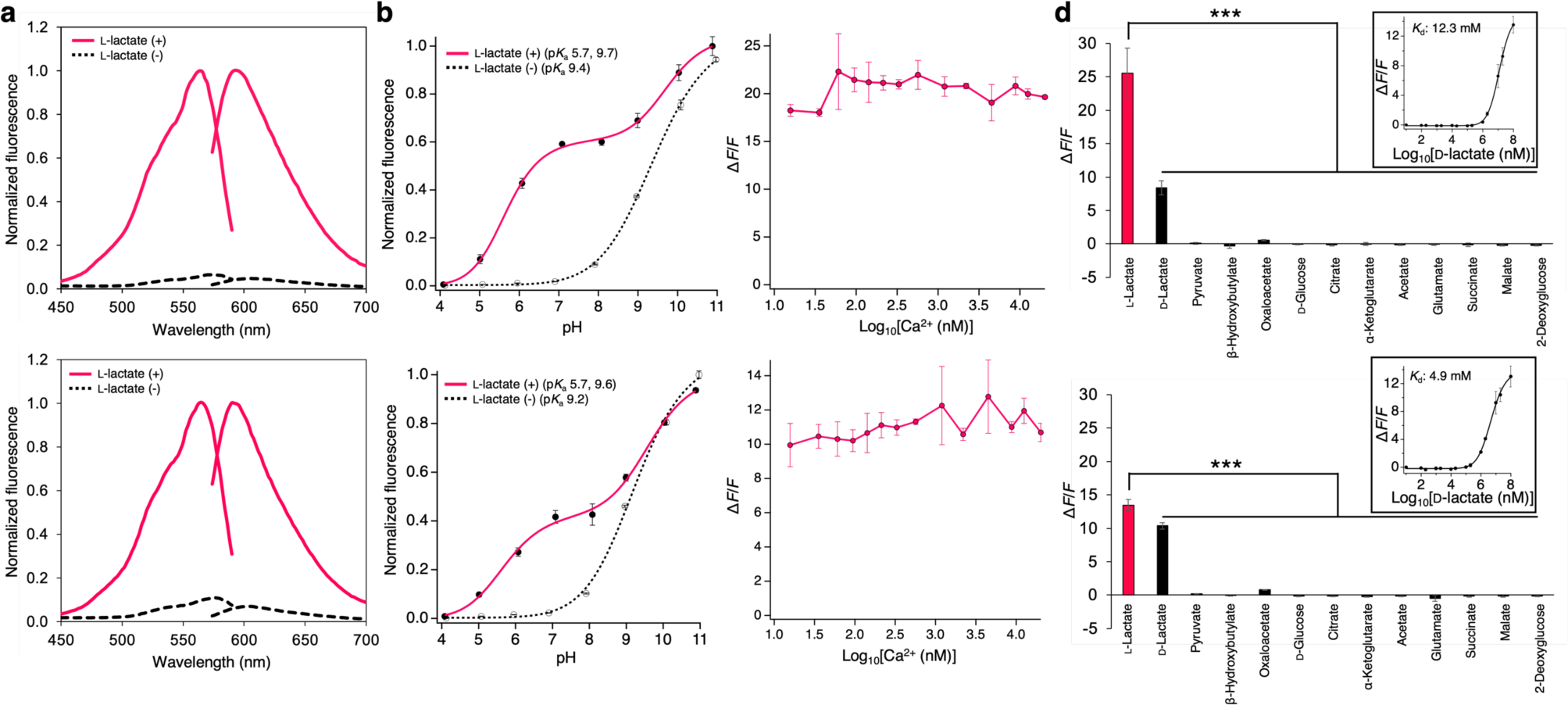
*In vitro* characterization of R-iLACCO1.1 and R-iLACCO1.2. (**a**) Excitation and emission spectra of R-iLACCO1.1 (upper) and R-iLACCO1.2 (bottom) in the presence (10 mM) and absence of l-Lactate. (**b**) pH titration curves of R-iLACCO1.1 (upper) and R-iLACCO1.2 (bottom) in the presence (100 mM) and absence of l-Lactate. *n* = 3 experimental triplicates (mean ± s.d.). (**c**) Δ*F*/*F* plot of R-iLACCO1.1 (upper) and R-iLACCO1.2 (bottom) as a function of Ca^2+^ in treatment with 100 mM l-Lactate. *n* = 3 experimental triplicates (mean ± s.d.). (**d**) Pharmacological specificity of R-iLACCO1.1 (upper) and R-iLACCO1.2 (bottom). Concentration of each metabolite is 10 mM. *n* = 3 experimental triplicates (mean ± s.d.). Statistical analyses were performed using one-way analysis of variance (ANOVA) with the Dunnett’s post hoc tests. \*\*\**p* < 0.0001.

**Supplementary Table 1.**
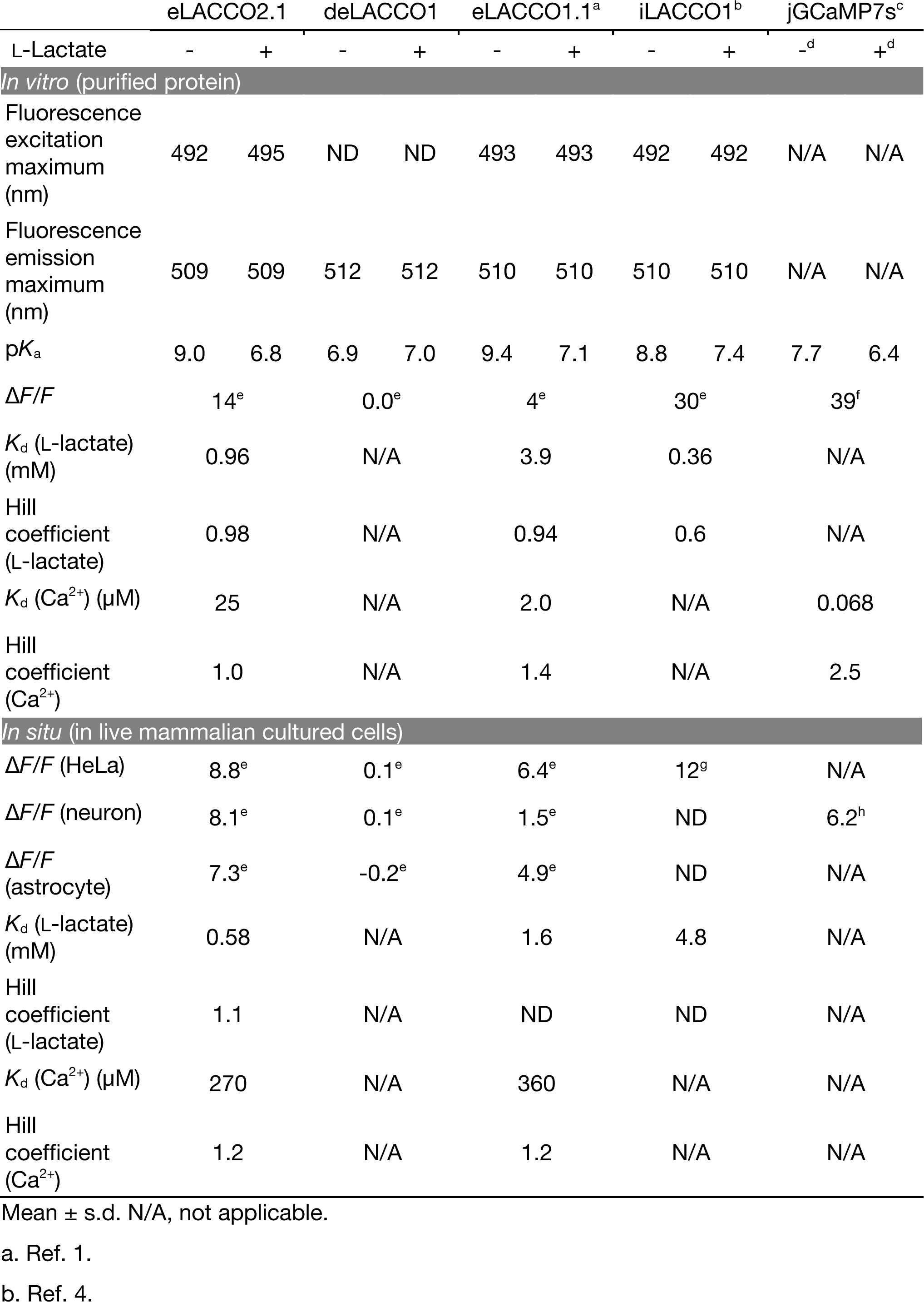
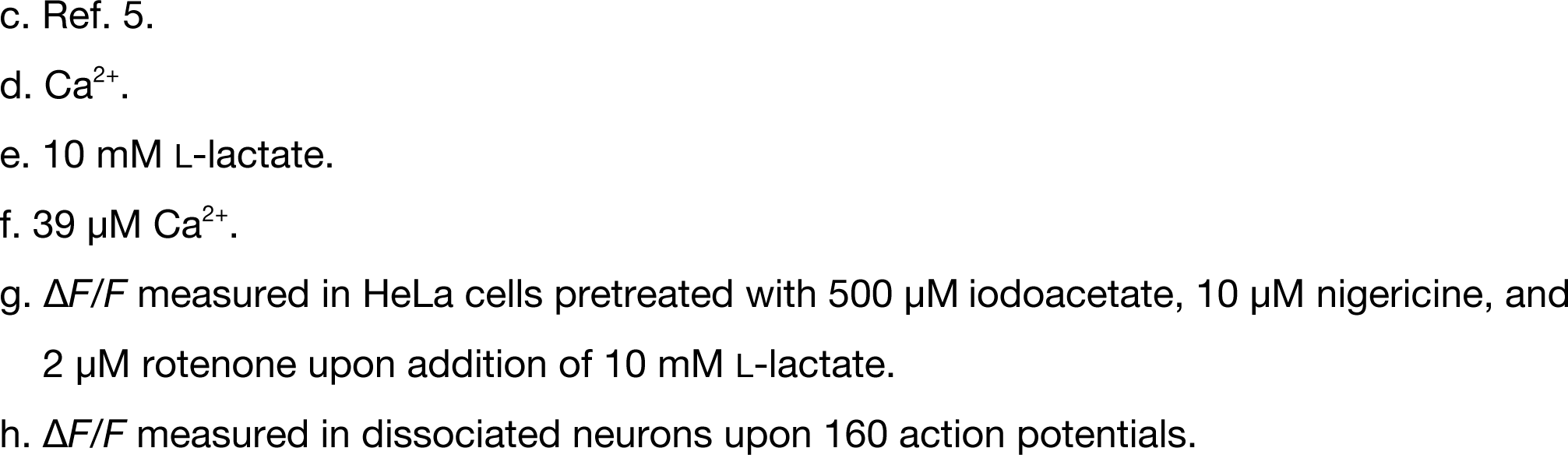
Biochemical parameters of eLACCO variants.

|  | eLACCO2.1 |  | deLACCO1 |  | eLACCO1.1<sup>a</sup> |  | iLACCO1<sup>b</sup> |  | jGCaMP7s<sup>c</sup> |  |
| --- | --- | --- | --- | --- | --- | --- | --- | --- | --- | --- |
| L-Lactate | - | + | - | + | - | + | - | + | -<sup>d</sup> | +<sup>d</sup> |
| *In vitro* (purified protein) |  |  |  |  |  |  |  |  |  |  |
| Fluorescence excitation maximum (nm) | 492 | 495 | ND | ND | 493 | 493 | 492 | 492 | N/A | N/A |
| Fluorescence emission maximum (nm) | 509 | 509 | 512 | 512 | 510 | 510 | 510 | 510 | N/A | N/A |
| pK<sub>a</sub> | 9.0 | 6.8 | 6.9 | 7.0 | 9.4 | 7.1 | 8.8 | 7.4 | 7.7 | 6.4 |
| $\Delta F/F$ | 14<sup>e</sup> | | 0.0<sup>e</sup> | | 4<sup>e</sup> | | 30<sup>e</sup> | | 39<sup>f</sup> | |
| K<sub>d</sub> (L-lactate) (mM) | 0.96 |  | N/A |  | 3.9 |  | 0.36 |  | N/A |  |
| Hill coefficient (L-lactate) | 0.98 |  | N/A |  | 0.94 |  | 0.6 |  | N/A |  |
| K<sub>d</sub> (Ca<sup>2+</sup>) (μM) | 25 |  | N/A |  | 2.0 |  | N/A |  | 0.068 |  |
| Hill coefficient (Ca<sup>2+</sup>) | 1.0 |  | N/A |  | 1.4 |  | N/A |  | 2.5 |  |
| *In situ* (in live mammalian cultured cells) |  |  |  |  |  |  |  |  |  |  |
| $\Delta F/F$ (HeLa) | 8.8<sup>e</sup> | | 0.1<sup>e</sup> | | 6.4<sup>e</sup> | | 12<sup>g</sup> | | N/A | |
| $\Delta F/F$ (neuron) | 8.1<sup>e</sup> | | 0.1<sup>e</sup> | | 1.5<sup>e</sup> | | ND | | 6.2<sup>h</sup> | |
| $\Delta F/F$ (astrocyte) | 7.3<sup>e</sup> | | -0.2<sup>e</sup> | | 4.9<sup>e</sup> | | ND | | N/A | |
| K<sub>d</sub> (L-lactate) (mM) | 0.58 |  | N/A |  | 1.6 |  | 4.8 |  | N/A |  |
| Hill coefficient (L-lactate) | 1.1 |  | N/A |  | ND |  | ND |  | N/A |  |
| K<sub>d</sub> (Ca<sup>2+</sup>) (μM) | 270 |  | N/A |  | 360 |  | N/A |  | N/A |  |
| Hill coefficient (Ca<sup>2+</sup>) | 1.2 |  | N/A |  | 1.2 |  | N/A |  | N/A |  |

Mean ± s.d. N/A, not applicable.
a. Ref. 1.
b. Ref. 4.
- c. Ref. 5. - d. $\text{Ca}^{2+}$ . - e. 10 mM L-lactate. - f. 39 $\mu\text{M}$ $\text{Ca}^{2+}$ . - g. $\Delta F/F$ measured in HeLa cells pretreated with 500 $\mu\text{M}$ iodoacetate, 10 $\mu\text{M}$ nigericine, and 2 $\mu\text{M}$ rotenone upon addition of 10 mM L-lactate. - h. $\Delta F/F$ measured in dissociated neurons upon 160 action potentials.

**Supplementary Table 2.**
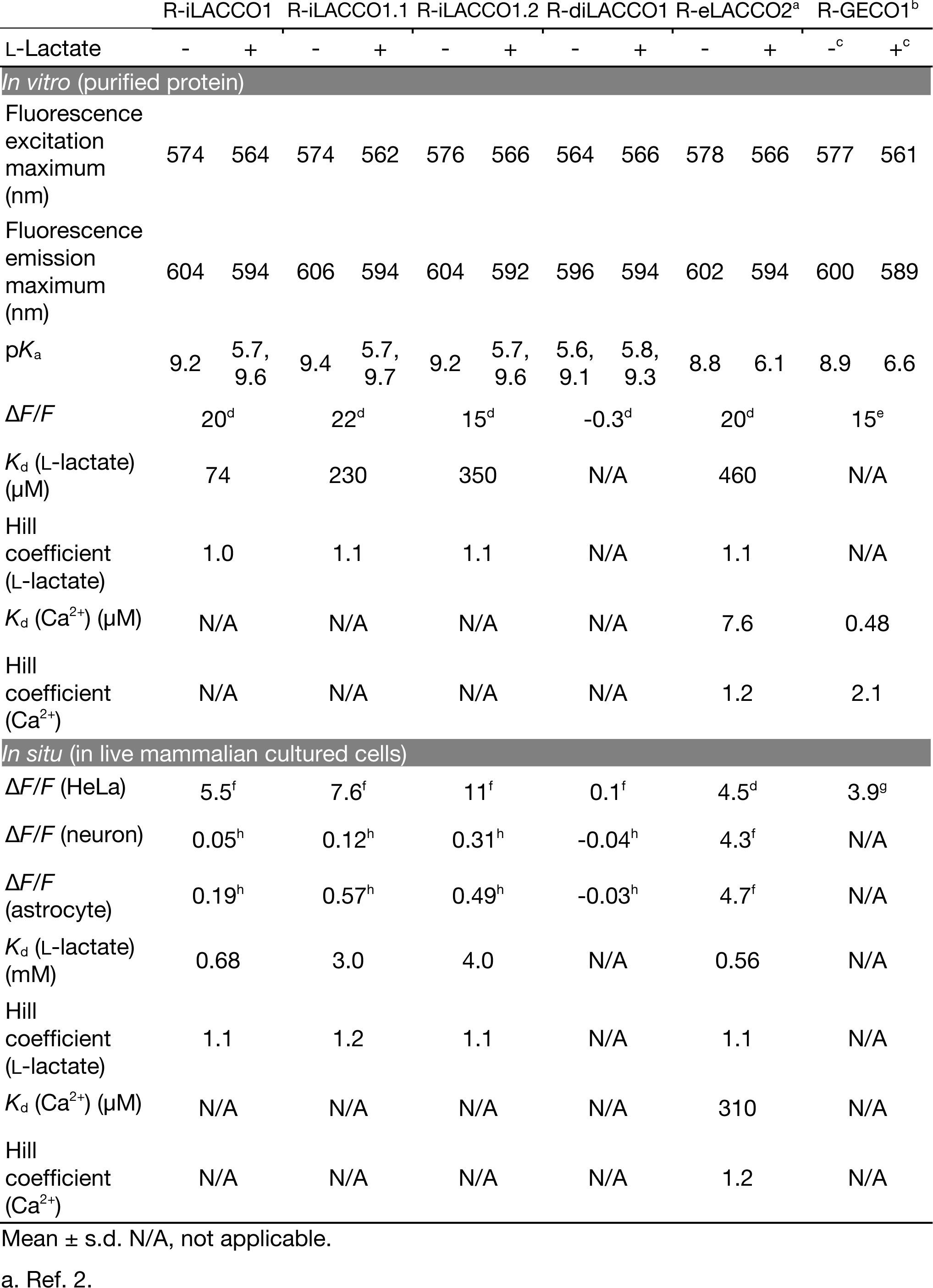
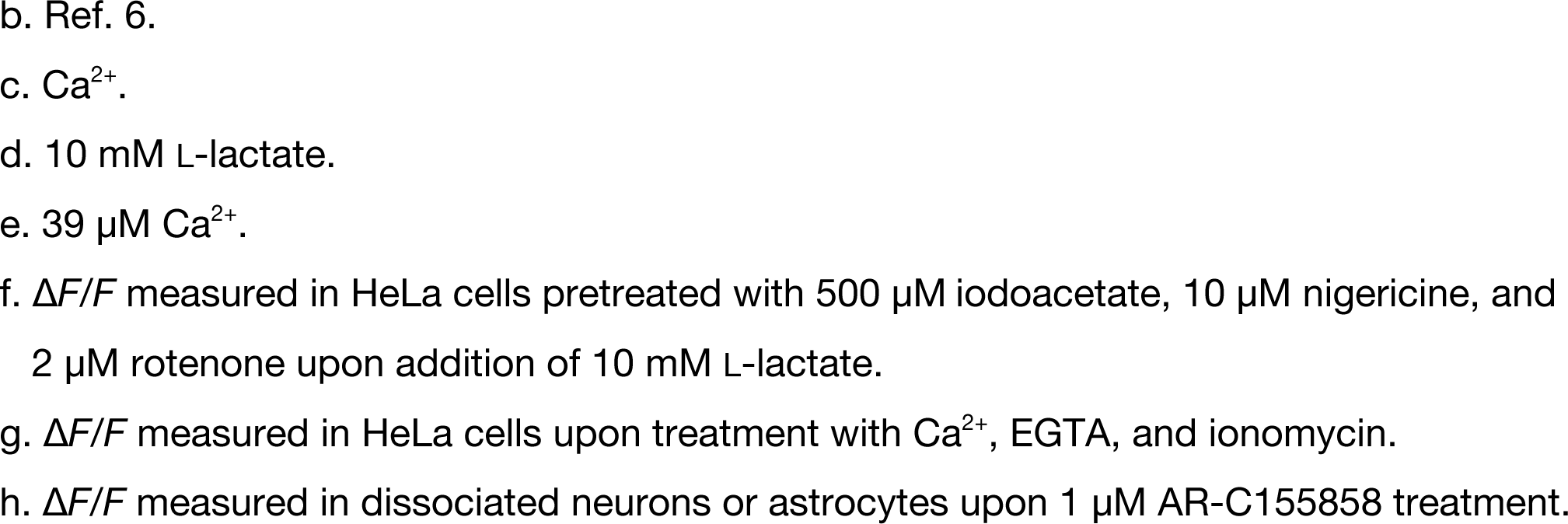
Biochemical parameters of R-iLACCO variants.

|  | R-iLACCO1 |  | R-iLACCO1.1 |  | R-iLACCO1.2 |  | R-diLACCO1 |  | R-eLACCO2<sup>a</sup> |  | R-GECO1<sup>b</sup> |  |
| --- | --- | --- | --- | --- | --- | --- | --- | --- | --- | --- | --- | --- |
| L-Lactate | - | + | - | + | - | + | - | + | - | + | -<sup>c</sup> | +<sup>c</sup> |
| *In vitro* (purified protein) |  |  |  |  |  |  |  |  |  |  |  |  |
| Fluorescence excitation maximum (nm) | 574 | 564 | 574 | 562 | 576 | 566 | 564 | 566 | 578 | 566 | 577 | 561 |
| Fluorescence emission maximum (nm) | 604 | 594 | 606 | 594 | 604 | 592 | 596 | 594 | 602 | 594 | 600 | 589 |
| pK<sub>a</sub> | 9.2 | 5.7, 9.6 | 9.4 | 5.7, 9.7 | 9.2 | 5.7, 9.6 | 5.6, 9.1 | 5.8, 9.3 | 8.8 | 6.1 | 8.9 | 6.6 |
| $\Delta F/F$ | 20<sup>d</sup> | | 22<sup>d</sup> | | 15<sup>d</sup> | | -0.3<sup>d</sup> | | 20<sup>d</sup> | | 15<sup>e</sup> | |
| K<sub>d</sub> (L-lactate) (μM) | 74 |  | 230 |  | 350 |  | N/A |  | 460 |  | N/A |  |
| Hill coefficient (L-lactate) | 1.0 |  | 1.1 |  | 1.1 |  | N/A |  | 1.1 |  | N/A |  |
| K<sub>d</sub> (Ca<sup>2+</sup>) (μM) | N/A |  | N/A |  | N/A |  | N/A |  | 7.6 |  | 0.48 |  |
| Hill coefficient (Ca<sup>2+</sup>) | N/A |  | N/A |  | N/A |  | N/A |  | 1.2 |  | 2.1 |  |
| *In situ* (in live mammalian cultured cells) |  |  |  |  |  |  |  |  |  |  |  |  |
| $\Delta F/F$ (HeLa) | 5.5<sup>f</sup> | | 7.6<sup>f</sup> | | 11<sup>f</sup> | | 0.1<sup>f</sup> | | 4.5<sup>d</sup> | | 3.9<sup>g</sup> | |
| $\Delta F/F$ (neuron) | 0.05<sup>h</sup> | | 0.12<sup>h</sup> | | 0.31<sup>h</sup> | | -0.04<sup>h</sup> | | 4.3<sup>f</sup> | | N/A | |
| $\Delta F/F$ (astrocyte) | 0.19<sup>h</sup> | | 0.57<sup>h</sup> | | 0.49<sup>h</sup> | | -0.03<sup>h</sup> | | 4.7<sup>f</sup> | | N/A | |
| K<sub>d</sub> (L-lactate) (mM) | 0.68 |  | 3.0 |  | 4.0 |  | N/A |  | 0.56 |  | N/A |  |
| Hill coefficient (L-lactate) | 1.1 |  | 1.2 |  | 1.1 |  | N/A |  | 1.1 |  | N/A |  |
| K<sub>d</sub> (Ca<sup>2+</sup>) (μM) | N/A |  | N/A |  | N/A |  | N/A |  | 310 |  | N/A |  |
| Hill coefficient (Ca<sup>2+</sup>) | N/A |  | N/A |  | N/A |  | N/A |  | 1.2 |  | N/A |  |

Mean ± s.d. N/A, not applicable.
a. Ref. 2.
- b. Ref. 6. - c. $\text{Ca}^{2+}$ . - d. 10 mM L-lactate. - e. 39 $\mu\text{M}$ $\text{Ca}^{2+}$ . - f. $\Delta F/F$ measured in HeLa cells pretreated with 500 $\mu\text{M}$ iodoacetate, 10 $\mu\text{M}$ nigericine, and 2 $\mu\text{M}$ rotenone upon addition of 10 mM L-lactate. - g. $\Delta F/F$ measured in HeLa cells upon treatment with $\text{Ca}^{2+}$ , EGTA, and ionomycin. - h. $\Delta F/F$ measured in dissociated neurons or astrocytes upon 1 $\mu\text{M}$ AR-C155858 treatment.

